# Simultaneous Targeting of KRAS and CDK4 Synergistically Suppresses Pancreatic Cancer Cells

**DOI:** 10.1101/2025.01.11.632518

**Authors:** Maj-Britt Paulsohn, Klara Henrike Frahnert, Xin Fang, Carolin Schneider, Constanza Tapia Contreras, Günter Schneider, Elisabeth Hessmann, Matthias Dobbelstein

## Abstract

Mutant Ras oncoproteins, particularly KRAS, are among the most prevalent drivers of cancer. Small-molecule inhibitors of KRAS have been developed, bearing high potential for cancer therapy but considerable risk of resistance development. To avoid cancer cell adaptation, effective combinatorial partners for increasing immediate efficacy remain to be explored. Here, we demonstrate that combining the KRAS inhibitor Sotorasib with the CDK4/6 inhibitor Palbociclib synergistically eliminates pancreatic ductal adenocarcinoma (PDAC) cells and organoids harboring KRAS G12C mutations. This synergy was particularly pronounced after drug washout, indicating a durable impact. Similar effects were observed in non-small-cell lung cancer (NSCLC) cells. Additionally, MRTX1133, a KRAS G12D inhibitor, synergized with Palbociclib to suppress KRAS G12D-mutant PDAC-derived cells. Mechanistically, these combinations induced sustained cell cycle arrest through reduced RB phosphorylation, decreased E2F1 levels, and increased CDKN1B/p27 expression. Deletion of CDKN1B largely rescued tumor cell proliferation, underscoring its critical role in mediating the observed synergy. These findings support the therapeutic potential of combining KRAS and CDK4/6 inhibitors for treating PDAC and other Ras-driven cancers.

## INTRODUCTION

Mutant Ras proteins are among the most common drivers of human cancer (Pylayeva-Gupta et al., 2011), a fact recognized for over four decades (Parada et al., 1982). Of the Ras protein family, KRAS is the most commonly mutated oncogene, particularly in pancreatic ductal adenocarcinoma (PDAC), non-small cell lung cancer (NSCLC), colorectal cancer and other malignancies (Singhal et al., 2024). Ras proteins function as GTP-binding proteins (G-proteins) that mediate receptor-driven signaling to intracellular kinases, notably through the RAF-MEK-ERK and PI3K-AKT-mTOR pathways. The majority of KRAS mutations occur at codon 12, with G12D, G12V, and G12C being the most prevalent (Lee et al., 2022). These mutations impair GTP hydrolysis, thereby prolonging signaling activity and driving oncogenesis.

Despite the pivotal role of KRAS mutations in cancer progression, the development of effective inhibitors has been challenging (Moore et al., 2020). Early attempts to inhibit KRAS through farnesylation interference were unsuccessful in clinical translation (Prendergast et al., 1996). However, significant breakthroughs were achieved in targeting mutant KRAS (Janes et al., 2018; Ostrem et al., 2013), which led to the introduction of Sotorasib, the first FDA-approved KRAS G12C inhibitor (Hong et al., 2020; Skoulidis et al., 2021). Sotorasib covalently binds to the mutant cysteine residue in KRAS G12C, effectively inhibiting its activity (Canon et al., 2019). Subsequent advancements led to the development of inhibitors targeting other KRAS mutations, such as G12D, offering new therapeutic opportunities for cancers like PDAC that are notoriously difficult to treat (Isermann et al., 2024; Punekar et al., 2022).

Despite these advances, resistance to KRAS inhibitors has emerged as a significant hurdle (Isermann et al., 2024). This highlights the urgent need for strategies that achieve sustainable elimination of KRAS-mutant cancer cells, through mechanisms such as inducing cell death or senescence, rather than transient cell cycle arrest. This challenge parallels earlier concerns with cyclin-dependent kinase 4 (CDK4) inhibitors, initially believed to induce only transient cell cycle arrest (Goel et al., 2018). CDK4 phosphorylates and inactivates the RB family of tumor suppressor proteins, which suppress E2F-driven transcription and proliferation. While CDK4 inhibitors were initially met with scepticism, they have since become a cornerstone of cancer therapy, particularly in breast cancer (Beaver et al., 2015; O’Leary et al., 2016), though their precise mechanisms of action remain incompletely understood (Watt and Goel, 2022).

The concept of synthetic lethality—combining targeted inhibitors to achieve permanent cancer cell elimination—was first proposed two decades ago (Kaelin, 2005). This approach holds particular promise for KRAS inhibitors, as effective combination therapies may overcome resistance and enhance therapeutic outcomes (Isermann et al., 2024; Perurena et al., 2024). Based on this rationale, we hypothesized that KRAS and CDK4 inhibitors could form a synthetic lethal combination capable of eradicating cancer cells. Supporting this idea, preclinical studies have shown that combining CDK4 inhibitors with agents targeting downstream signaling pathways of KRAS effectively suppresses tumor growth in PDAC models (Cheng et al., 2024; Goodwin et al., 2023; Willobee et al., 2021). With the recent availability of direct KRAS inhibitors, it becomes critical to investigate whether their combination with CDK4 inhibitors could produce synergistic and durable therapeutic effects. Here, we demonstrate that the combination of KRAS and CDK4 inhibitors synergistically and sustainably suppresses PDAC and NSCLC cell proliferation and viability across multiple experimental systems, including cell lines and organoids. This combination therapy significantly enhances the prospects for treating PDAC and other challenging cancers that remain difficult to cure.

## RESULTS

### The KRAS mutant G12C-targeting drug Sotorasib synergizes with Palbociclib to suppress the growth of pancreatic cancer cells and organoids, with sustainability after drug removal

Previous studies pointed to CDK4 as a promising combination partner for KRAS inhibitors to eliminate PDAC cells. This became evident when we re-analysed a published CRISPR interference (CRISPRi) screen of MIA PaCa-2 cells treated with the KRAS G12C inhibitor ARS-1620 (Lou et al., 2019). The analysis revealed that the deletion of cyclin D1 (CCND1) or its interaction partner CDK4 each conferred some of the strongest sensitization towards the drug (Suppl. Figure 1A). Consistent with these findings, combining inhibitors of MEK/MAPK2 (Cheng et al., 2024; Willobee et al., 2021) or ERK/MAPK3 (Adamia et al., 2022; Goodwin et al., 2023; Jagirdar et al., 2024) with CDK4 inhibition has been shown to effectively eliminate cancer cells of entities such as PDAC or multiple myeloma. Based on this evidence, we investigated whether the simultaneous inhibition of KRAS and CDK4 could synergistically suppress PDAC cell proliferation. Initially, we picked the most established KRAS inhibitor, Sotorasib (also known as AMG510), which specifically targets the G12C mutant of KRAS through covalent modification (Canon et al., 2019; Ostrem et al., 2013; Pantsar, 2020). Indeed, treating MIA PaCa-2 cells with Sotorasib along with Palbociclib profoundly suppressed cell confluence as well as viability (Figure 1A, Suppl. Figure 1B). Notably, the synergy was even more pronounced four days after drug removal, indicating sustainable efficacy of the drug combination. This was confirmed by the BLISS synergy score. Similar observations were made using the cell line 51T-2D, which we had established from a patient-derived organoid (PDO) of PDAC (Tapia Contreras et al., 2024). Like MIA PaCa-2 cells, 51T-2D cells carry the KRAS G12C mutation, which is otherwise rare in PDAC, thus providing an additional opportunity to test the impact of Sotorasib and its cooperation with Palbociclib. Here again, we observed sustained growth arrest and drug synergy (Figure 1B, Suppl. Figure 1C). Like Palbociclib, the CDK4 inhibitor Abemaciclib diminished cell proliferation and viability in combination with Sotorasib in MIA PaCa-2 and 51T-2D cells (Suppl. Figure 1D, E, F, G). In contrast, and as expected, Sotorasib did not modulate the impact of Palbociclib on KPC cells, which carry the KRAS G12D mutation (Suppl. Figure 1H). In Sotorasib-susceptible cells, however, the drug combination diminished the proportion of cells undergoing DNA replication, as revealed by incorporation of a nucleoside analogue (Suppl. Figure 1I, J). The drugs also enhanced the fraction of dead cells, albeit on a low overall level, as assayed by annexin V staining, the uptake of propidium iodide, and PARP1 cleavage (Suppl. Figure 1K, L). We also combined Palbociclib with inhibitors that target signaling factors downstream of KRAS, i.e. MEK or ERK. Such inhibitors also displayed synergy with the CDK4 inhibitor, albeit to a lesser degree than KRAS inhibitors in the investigated experimental settings (Suppl. Figure 1M, N).

**Figure 1:**
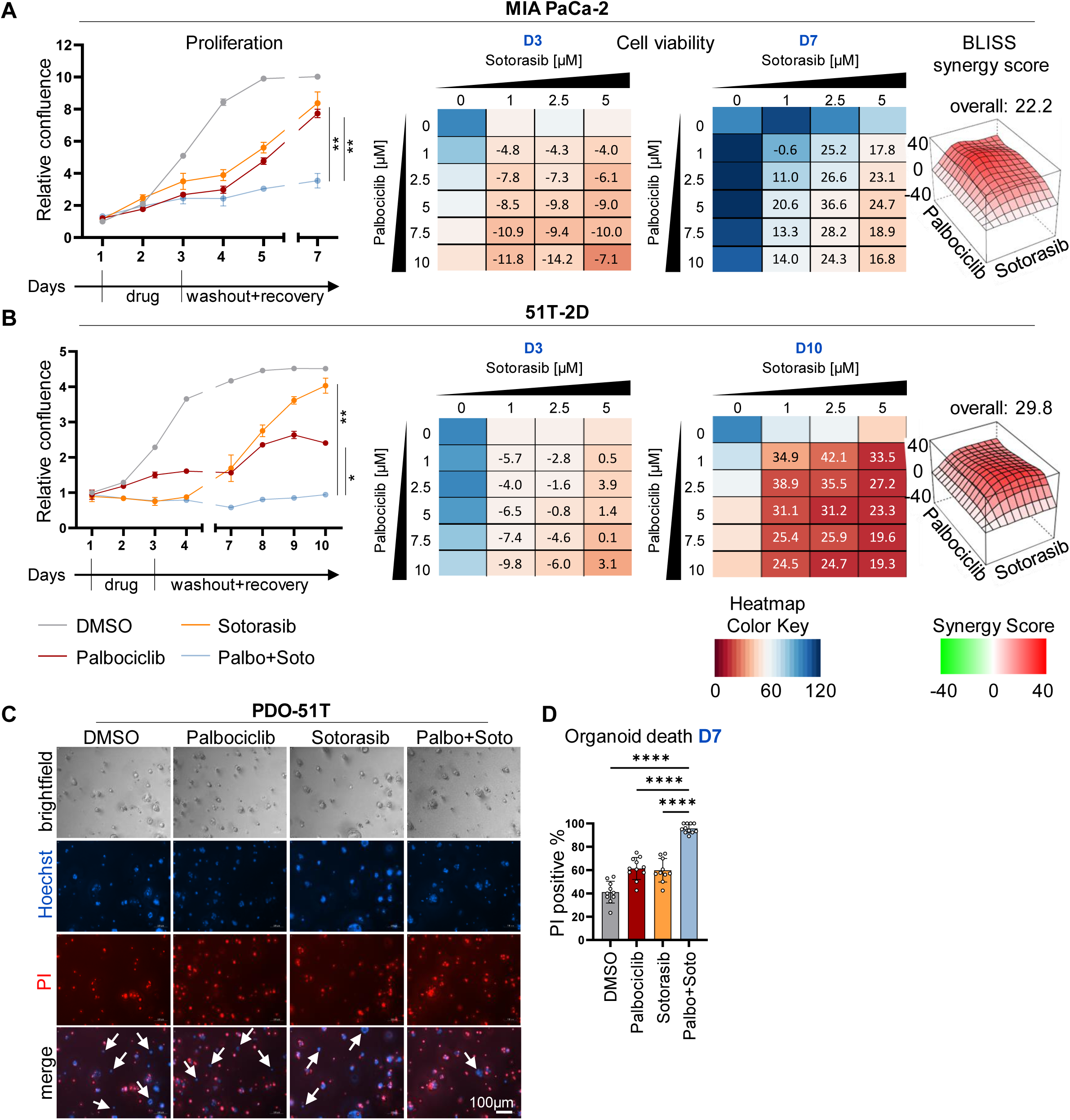
The KRAS mutant G12C-targeting drug Sotorasib synergizes with Palbociclib to suppress the growth of pancreatic cancer cells and organoids, with sustainability after drug removal. (A) (left) Proliferation of MIA PaCa-2 cells, determined by automated microscopy (Celigo®) upon treatment with DMSO, 5 µM Palbociclib, 2.5 µM Sotorasib, or the combination. Cells were treated for 48 h, followed by four days of recovery in the absence of drugs. The graphs indicate the means of three technical replicates ± SD. (right) Heat maps reflect the viability of MIA PaCa-2 cells right after treatment (D3) and after recovery (D7), normalized to the DMSO controls. Bliss synergy score values were determined based on cell viability. Drug values >10 reflect synergy, whereas values between -10 and 10 correspond to an additive mechanism. The BLISS synergy map of D7 is depicted. (B) (left) Proliferation of 51T-2D cells upon treatment as in (A) (right) Heat maps show the viability of 51T-2D cells right after treatment (D3) and after recovery (D10), normalized to the DMSO control. BLISS synergy map of D10. (C) PDO-51T organoids (KRAS G12C) treated with 5 µM Palbociclib and 2.5 µM Sotorasib for 48 h, followed by four days of recovery without drugs. Representative brightfield images of day 7 (D7) as well as Hoechst 33342 and PI staining are presented. Scale bar, 100 µm. (D) Hoechst 33342 and PI staining was used to quantify the viability of organoids at day 7 (D7). Statistical analyses: A, B unpaired t-test (A, B: of AUC); D, one-way ANOVA followed by Tukey’s multiple comparison; ns: not significant, *p ≤ 0.05, **p ≤ 0.01, ***p ≤ 0.001, ****p ≤ 0.0001.

Since tumor cell organoids often reflect the behaviour of actual tumors more closely than 2D-cultures, we also investigated the impact of Sotorasib and Palbociclib on the survival of PDO-51T. Indeed, the drugs cooperated to enhance the uptake of propidium iodide, indicating enhanced death of organoid cells (Figure 1C, D). We conclude that the combination of Sotorasib and a CDK4 inhibitor strongly synergizes to compromise the proliferation and viability of PDAC cells and organoids.

### Palbociclib and KRAS inhibitors synergize to suppress PDAC and NSCLC cell proliferation, and to induce senescence

Sotorasib exclusively targets the KRAS mutant G12C, which is rare in PDAC. To raise a perspective for broader treatment options, we evaluated a similar approach with additional pancreatic cancer cells that carry the more frequently encountered mutation G12D in KRAS. At present, inhibitors of KRAS G12D have not yet been approved for clinical use. However, the KRAS G12D inhibitor MRTX1133 (Hallin et al., 2022; Wang et al., 2022) is currently tested in phase I and II trials for advanced solid tumors (NCT05737706). We evaluated the effect of inhibiting KRAS G12D and CDK4, alone or in combination. MRTX1133 and Palbociclib diminished the proliferation and the viability of human AsPC-1 and murine KPC cells (Figure 2A, B, Suppl. Figure 2A, B). Synergy was revealed by the Bliss score, in particular seven days after drug removal. In addition, we analysed cell lines established from patient-derived PDAC xenografts (GöCDX7, GöCDX52 and GöCDX53, all KRAS G12D). GöCDX52 cells responded synergistically towards Palbociclib and MRTX1133, with significantly impaired cell proliferation and viability, again particularly seven days after drug removal (Suppl. Figure 2 E). GöCDX7 and GöCDX53 showed reduced growth paired with synergistic effects on viability, and this was confirmed by the BLISS score, too (Suppl. Figure 2F, G). Again, these results suggest that combining inhibitors of KRAS and CDK4 represents a synergistic and sustainable strategy to treat KRAS G12D mutant PDAC.

**Figure 2:**
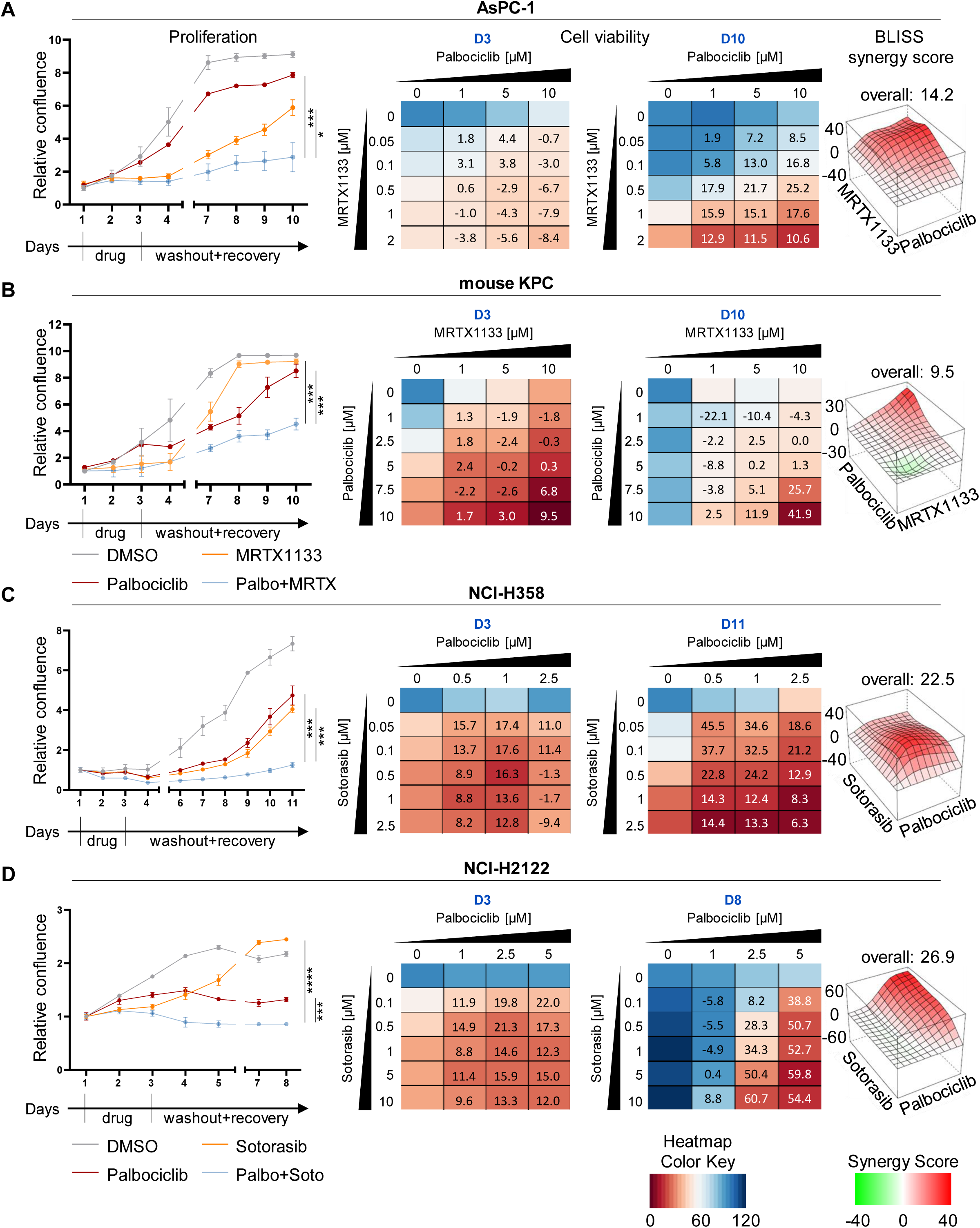
Palbociclib and KRAS inhibitors synergize to suppress PDAC and NSCLC cell proliferation, and to induce senescence. (A-D) All cells were treated and observed as in Fig. 1, with the following specifics: (A) AsPC-1 cells (human PDAC), 5 µM Palbociclib and 0.5 µM MRTX1133, measurements at D3 and D10. (B) KPC cells (murine PDAC), 5 µM Palbociclib, 5 µM MRTX1133, evaluation at D3 and D10. (C) NCl-H358 cells (human NSCLC), 0.5 µM Palbociclib, 50 nM Sotorasib, evaluation at D3 and D10. (D) NCl-H2122 (human NSCLC), 2.5 µM Palbociclib, 5 µM Sotorasib, evaluation at D3 and D8. Statistical analyses: A, B, C, D unpaired t-test (A, B, C, D: of AUC); ns: not significant, *p ≤ 0.05, **p ≤ 0.01, ***p ≤ 0.001, ****p ≤ 0.0001.

Besides PDAC, NSCLC is another entity with a high proportion of KRAS mutations. Here, the mutation G12C is most prominent, justifying Sotorasib monotherapy according to a recent study (Skoulidis et al., 2021). However, when treating the NSCLC-derived cell lines NCl-H358 and NCl-H2122 with Sotorasib, the cells were transiently stalled but resumed proliferation within a few days; in contrast, additional CDK4 inhibition largely prevented cell outgrowth (Figure 2C, D, Suppl. Figure 2C, D). Once again, we observed that the combination of KRAS and CDK4 inhibition strongly diminished cell proliferation and viability. Synergistic effects were observed already after 48 h of treatment, and this synergy became more robust after the additional recovery period. In summary, we provide evidence of sustainable and synergistic growth arrest by inhibiting CDK4 and KRAS G12C/D in PDAC and NSCLC cells.

In AsPC-1 cells, the drugs Palbociclib and MRTX1133 not only gave rise to ceased proliferation, but also to cell senescence, as revealed by senescence-associated beta-galactosidase (SAB) staining. Here again, the combination of the drugs yielded more pronounced effects than either drug alone, after 24 h treatment and after the 48 h recovery period (Suppl. Figure 2H, I). Similar observation of senescence were made using Sotorasib together with Palbociclib to treat NCl-H358 cells (Suppl. Figure 2J, K). Thus, the drug combinations induce senescence, providing one mechanistic clue to explain their activities against cancer cells.

### Inhibitors of KRAS G12C and CDK4 cooperate to switch the transcriptome towards ceased proliferation

In search for mechanisms that underlie the observed synergy between inhibitors of KRAS G12C and CDK4, we analysed mRNA levels in 51T-2D cells by deep sequencing, both immediately after treatment (Day3), and four days after washing off the drugs (Day7; PCA: Suppl. Figure 3A, B, Suppl. Table 1). For day3 the heat map of the z-scores depicts profound changes in gene expression between Palbociclib, Sotorasib and the combination treatment (Figure 3A). Differentially expressed genes upon combination treatment were correlated with the Molecular Signature Database (MSigDB). This revealed negative regulation of genes corresponding to the signatures of E2F targets, the G2M checkpoint, mTORC1 signaling as well as MYC targets (Figure 3B). Moreover, analysis of associated transcription factors pointed out E2F1, E2F4, MYC and FOXM1 (Figure 3C). Gene set enrichment analyses (GSEA) identified similar pathways in the comparison between the single use of either Palbociclib or Sotorasib with the combination of both drugs (Figure 3D). In addition, we found genes related to glycolysis and oxidative phosphorylation to be suppressed upon combination treatment. Normalized counts of E2F targets were significantly diminished, immediately after treatment and also after four days of washout (Figure 3E, Suppl. Figure 3C). Quantitative RT-PCR analyses confirmed that both inhibitors suppressed genes that need to be activated before the cell passes through the cell cycle checkpoints G1→S or G2→M, e.g. *CCNE1*/Cyclin E1 (Figure 3F), *CCND1*/Cyclin D1, *BIRC5*/Survivin, *CCNB1*/Cyclin B1, and *PLK4*. Importantly, when combining both drugs, the suppression of these genes was largely maintained even 48 h or four days after drug removal. In contrast, the single drugs, when applied to the cells and then removed, only transiently diminished the expression of the same genes and allowed their re-appearance after washout. These observations were made in 51T-2D (Suppl. Figure 3D, E), MIA PaCa-2 (Suppl. Figure 3F) and NCl-H358 cells (Suppl. Figure 3G). We conclude that the combined inhibition of KRAS and CDK4 sustainably suppresses the expression of genes that would be necessary to resume cell proliferation.

**Figure 3:**
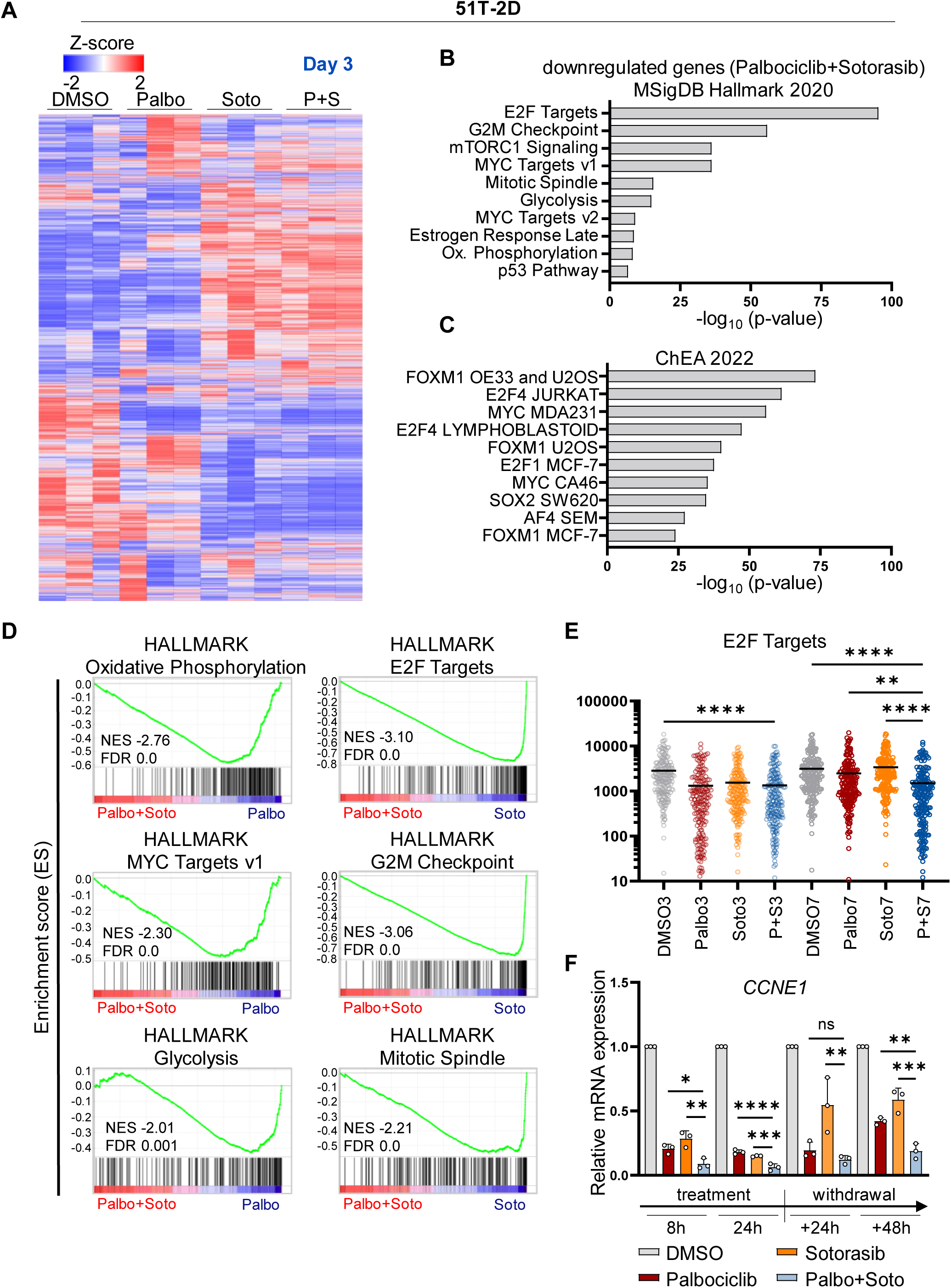
Inhibitors of KRAS G12C and CDK4 cooperate to switch the transcriptome towards ceased proliferation. (A) Heat map depicting differentially expressed (DE) genes according to the z-score after performing DeSeq2 analysis of four different samples (DMSO, 5 µM Palbociclib, 2.5 µM Sotorasib or combination treatment for 48 h (Day3), n=3) in 51T-2D cells. Only genes with |log2fold| ≥ 0.6, adjusted p-value (padj.) < 0.05, and baseMean ≥ 15 were included in the analysis. Suppl. Table 1 contains DE genes and normalized read counts. (B) Negatively regulated genes in response to Palbociclib + Sotorasib (48 h treatment) vs DMSO were correlated with the Molecular Signature Database (MSigDB) Hallmark 2020 and ChEA 2022 (C) dataset using the Enrichr platform to identify potentially impaired pathways. Top 10 (ChEA 2022: human only), p-value ranked (-log10). (D) Gene set enrichment analysis (GSEA) of combination treatment vs Palbociclib or Sotorasib monotreatment, after 48 h of treatment, determining hallmarks (h.all.v2023.2). (E) Normalised counts of E2F targets upon 48 h treatment (3) or 48 h treatment + four days of drug withdrawal (7). (F) Expression of E2F target gene *CCNE1* in 51T-2D cells treated with 5 µM Palbociclib, 2.5 µM Sotorasib or the combination, for 8 and 24 h, and drug withdrawal for 24 or 48 h. Data was normalized to *36B4* mRNA and is shown as mean ± SD. Statistical analyses: E, F one-way ANOVA followed by Tukey’s multiple comparison; ns: not significant, *p ≤ 0.05, **p ≤ 0.01, ***p ≤ 0.001, ****p ≤ 0.0001. Complete statistics in Suppl. Figure 3 C.

Similarly to the transcriptome analysis immediately after treatment, we performed RNA sequencing after four days of an additional washout phase and displayed the z-scores to compare expression levels (Suppl. Figure 3H). Here, we found further evidence for a permanent cell cycle arrest in the maintained suppression of gene sets with E2F targets and the G2M checkpoint (Suppl. Figure 3I, J). Cells that had been treated individually with Palbociclib or Sotorasib re-expressed E2F target genes; Sotorasib-treatment with washout resulted in even stronger expression of these genes than control treatment. In contrast, GSEA of cells receiving the combination treatment continuously repressed genes of the categories E2F target, G2M checkpoint or MYC target (Suppl. Figure 3K). All these gene sets and transcription factors are indispensable for cell cycle progression, corroborating our observations of permanent cell cycle arrest induced by the drug combination.

### Transcriptome alterations in PDAC cells treated with Palbociclib and MRTX1133

To further extend our mechanistic understanding of the cooperative inhibition of CDK4 and KRAS in PDAC cells, we analysed the transcriptome of AsPC-1 cells treated for 24 h with Palbociclib and the KRAS G12D inhibitor MRTX1133 (PCA: Suppl. Figure 4A, Suppl. Table 2). The heat map of the z-scores displays similarities between control and Palbociclib on the one hand and MRTX1133 and the combined treatment on the other hand (Figure 4A). Genes suppressed by both drugs correlate with E2F targets, G2M checkpoint, MYC targets, and mTORC1 signaling, based on MSigDB (Figure 4B). Similar to the transcriptome analysis obtained with Palbociclib and Sotorasib (Figure 3), the combination of Palbociclib with MRTX1133 revealed transcription factors E2F1, E2F4, MYC and FOXM1 to be associated with genes that were suppressed by the combination (Figure 4C). In addition, GSEA found MYC targets, E2F targets, G2M checkpoint and mTORC1 signaling negatively enriched in cells treated with the combination (Figure 4D). This is flanked by reduced normalized counts of E2F targets when combining Palbociclib and MRTX1133 (Figure 4E, Suppl. Figure 4B). Once again, we performed quantitative RT-PCR analyses to confirm that both drugs significantly impair the expression of genes required for the G1→S or G2→M transitions, e.g. *CCNE1*/Cyclin E1 (Figure 4F). While single drugs only transiently repressed these genes, the combination diminished the expression of genes such as *BIRC5*/Survivin, *BRCA1* and *CCNB1*/Cyclin B1 even after 48 h of drug withdrawal (Suppl. Figure 4C). Interestingly, genes of the CDKN2 family of CDK inhibitors were expressed more strongly immediately after treatment with Palbociclib and MRTX1133 (Suppl. Figure 4D), while in Palbociclib and Sotorasib treated 51T-2D cells the CDKN1 family genes were enriched (Suppl. Figure 3L), perhaps providing additional growth-inhibitory mechanisms. In conclusion, we find Palbociclib and MRTX1133 to suppress genes required to overcome cell cycle checkpoints and for continuous proliferation.

**Figure 4:**
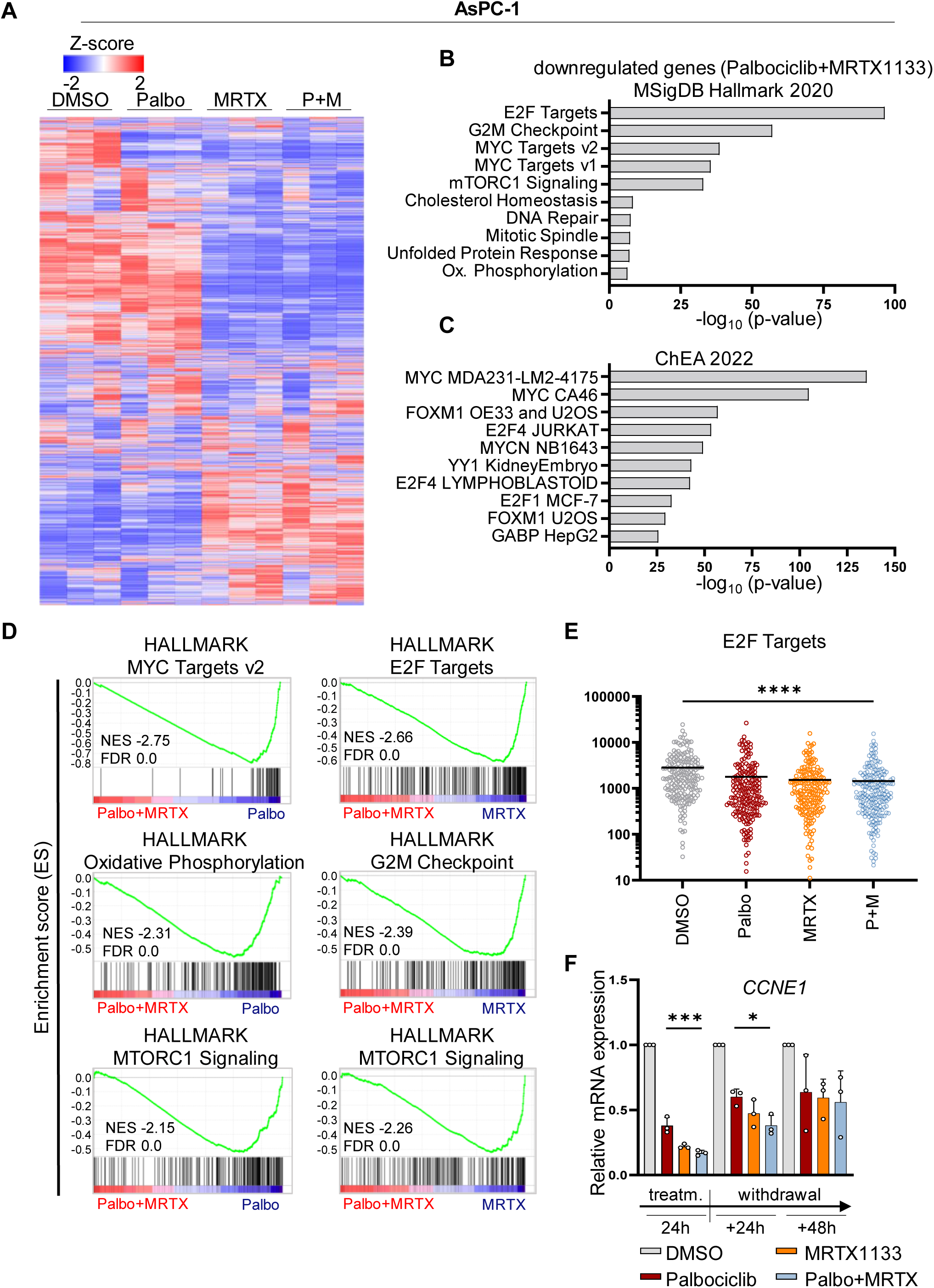
Transcriptome alterations in PDAC cells treated with Palbociclib and MRTX1133. (A) Heat map depicting DE genes according to the z-score after performing DeSeq2 analysis of four different samples (DMSO, 5 µM Palbociclib, 0.5 µM MRTX1133, or combination treatment, for 24 h (Day2), n=3) in AsPC-1 cells. Only genes with |log2fold| ≥ 0.6, adjusted p-value (padj.) < 0.05, and baseMean ≥ 15 were included in the analysis. Suppl. Table 2 contains DE genes and normalized read counts. (B) Downregulated genes in Palbociclib + MRTX1133 (24 h treatment) vs DMSO were correlated with the Molecular Signature Database (MSigDB) Hallmark 2020 and ChEA 2022 (C) datasets, using the Enrichr platform to identify potentially impaired pathways. Top 10 (ChEA 2022: human only), p-value ranked (-log10). (D) Gene set enrichment analysis (GSEA) of combination treatment vs Palbociclib or MRTX1133 monotherapy after 24 h of treatment; hallmarks (h.all.v2023.2). (E) Normalised counts of E2F targets upon 24 h treatment. (F) Expression of the E2F target gene *CCNE1* in AsPC-1 cells treated with 5 µM Palbociclib, 0.5 µM MRTX1133, or the combination, for 24 h, and drug withdrawal for 24 or 48 h. mRNA levels were normalized to *36B4* mRNA, mean ± SD. Statistical analyses: E, F one-way ANOVA followed by Tukey’s multiple comparison; ns: not significant, *p ≤ 0.05, **p ≤ 0.01, ***p ≤ 0.001, ****p ≤ 0.0001. Complete statistics in Suppl. Figure 4 B.

### Upon inhibition of KRAS and CDK4, E2F1 largely disappears while CDKN1B/p27 levels increase and RB family proteins adopt a hypophosphorylated state

To further understand how the two drugs achieve sustainable proliferation arrest, we assessed protein levels and phosphorylation states by immunoblot analyses. Immediately upon treatment, both drugs suppressed the phosphorylations that their targets or downstream signaling mediators would otherwise confer: ERK and AKT phosphorylation was diminished by Sotorasib, and RB1 phosphorylation by Palbociclib, in 51T-2D and MIA PaCa-2 cells (Suppl. Figure 5A, B). After drug removal for another 24 or 48 h, however, the drug combination had a more sustainable impact on cell cycle regulators than the single drugs. In particular, the phosphorylation of RB1 was still reduced when both drugs were combined, and the overall levels of RB1 and RBL1/p107 were also suppressed. In contrast, RBL2/p130 was augmented under the same conditions (Figure 5A, B). Remarkably, RBL2 is a key member of the repressive dimerization partner, RB-like, E2F and multi-vulval class B (DREAM) complex that represses the transcription of genes such as *BIRC5*, *BRCA1*, *CCNB1*, and *PLK4* (Engeland, 2018; Fischer et al., 2016), i.e. the same proliferation-associated genes that we had found suppressed by the drug combination (Suppl. Figures 3D, E, F, G, 4C). Similar observations were made in NCl-H358 cells (NSCLC; Suppl. Figure 5C) or when using the KRAS G12D antagonist MRTX1133 together with Palbociclib in AsPC-1 cells (PDAC; Suppl. Figure 5D). Furthermore, our analyses revealed that the levels of the key activating member of the E2F transcription factor family, E2F1, were reduced upon Palbociclib treatment, but even more strongly upon combined drug treatment (Figure 5C). In particular, four days after drug removal, E2F1 levels remained low upon treatment with the combination, while E2F1 was re-synthesized upon monotherapy. E2F1 is a known activator of the above-named genes (Rouillard et al., 2016). In addition, the mRNA encoding the cell cycle regulator CDKN1B/p27 was upregulated upon Sotorasib and combined treatment (Suppl. Figure 3D, E, F, G, 4C). However, the p27 protein levels appeared more strongly elevated upon combination treatment for 48 h (Figure 5D). In summary, these analyses elucidate the impact of the drug combinations on the transcriptome as follows: the combined reduction in E2F1 levels and the concurrent increase in the DREAM complex ensure sustained suppression of genes essential for bypassing cell cycle checkpoints. Additionally, the elevated levels of the cell cycle inhibitor p27 reinforce this effect. Consequently, the cells remain arrested in response to the drug combination treatment.

**Figure 5:**
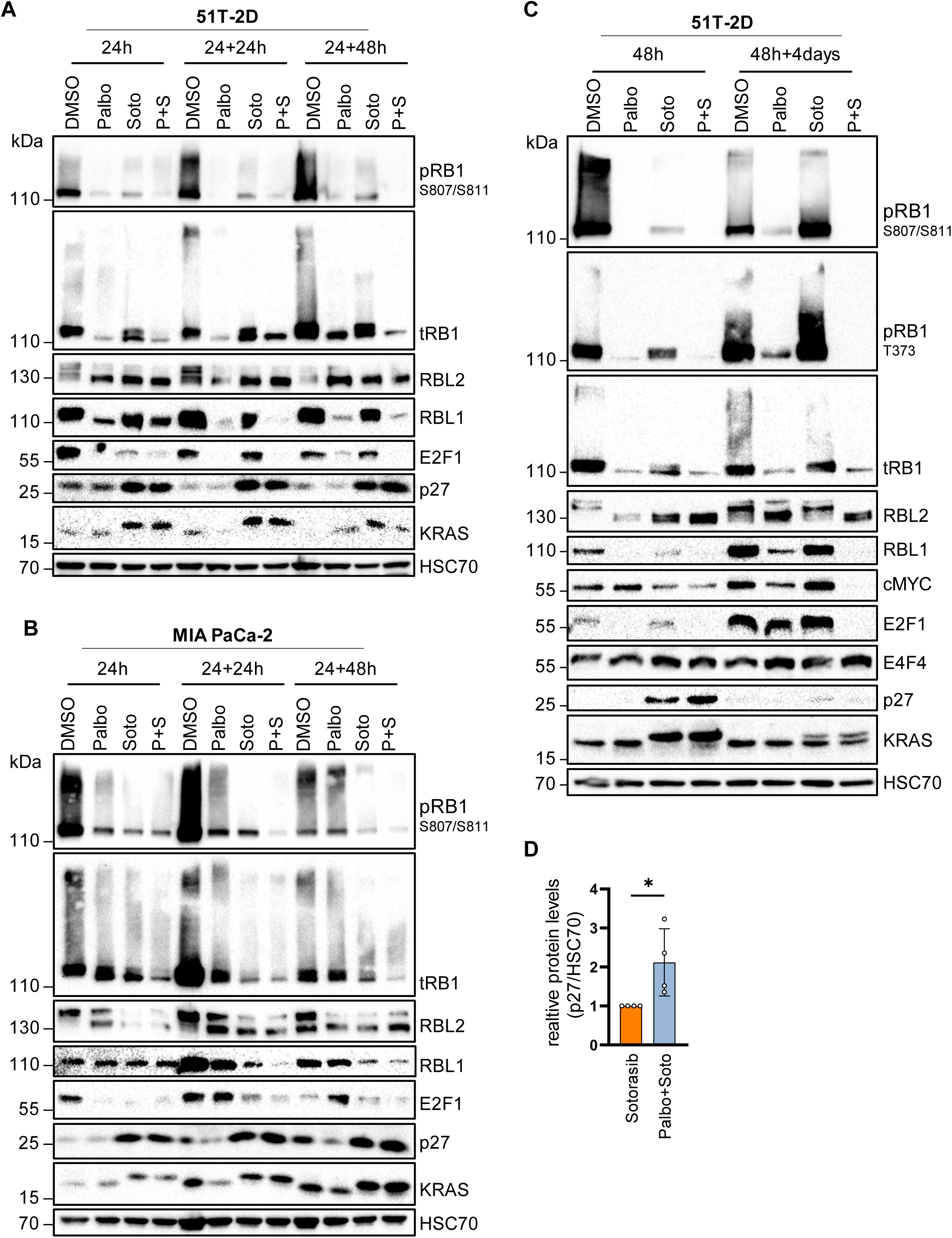
Upon inhibition of KRAS and CDK4, E2F1 largely disappears while CDKN1B/p27 levels increase and RB family proteins adopt a hypophosphorylated state. (A) Immunoblot analysis of whole-cell lysates of 51T-2D cells treated with DMSO, 5 µM Palbociclib, 2.5 µM Sotorasib, or both, for 24 h and wash-out for 24 or 48 h. HSC70 was detected as sample control. One representative immunoblot is shown; n=2. Sotorasib covalently binds to KRAS, reflected by an electrophoretic mobility shift. (B) Immunoblot analysis as in (A), using MIA PaCa-2 cells treated with DMSO, 10 µM Palbociclib, 5 µM Sotorasib or the combination; n=2. (C) Immunoblot analysis 51T-2D cells treated as in (A) for 48 h or 48 h + four days of drug withdrawal; n=2. (D) Quantification of p27 protein levels in relation to loading control (HSC70) corresponding to (C); n=4. Statistical analyses: D, unpaired t-test; ns: not significant, *p ≤ 0.05, **p ≤ 0.01, ***p ≤ 0.001, ****p ≤ 0.0001.

### Growth suppression by inhibitors of KRAS and CDK4 depends on the RB family proteins and p27

To investigate the mechanistic role of RB family proteins and p27 in the response to the drugs, we performed knockdown experiments by transfection of MIA PaCa-2 cells with siRNA. Depletion of RB1 significantly restored cell proliferation despite treatment with the drug combination (Figure 6A, Suppl. Figure 6A). While RBL1 knockdown merely attenuated the response to Sotorasib (Figure 6B), RBL2 depletion yielded some recovery of cells that were treated with the combination (Figure 6C). E2F4 depletion significantly accelerated cell recovery from Sotorasib treatment (Figure 6D) but only weakly rescued the cells from combined treatment. FOS depletion significantly diminished the proliferation of cells upon treatment with Palbociclib or Sotorasib (Figure 6E). With CDKN1B knockdown, Sotorasib-treated cells resumed proliferation rapidly, whereas the response to Palbociclib monotherapy was only slightly affected (Figure 6F, G, H). Remarkably, however, CDKN1B depletion allowed significant recovery upon combination treatment, especially when using low doses of Palbociclib. All knockdowns were verified by immunoblot and/or quantitative RT-PCR (Suppl. Figure 6 B, C, D). To further corroborate these results, we deleted CDKN1B in the murine PDAC cell line 8661 (knockout, KO; Suppl. Figure 6E). Cells with CDKN1B KO were less susceptible to either KRAS or CDK4 inhibition, compared to parental wild type (WT) cells. Most notably, CDKN1B KO cells recovered from the drug combination almost completely, while WT cells continued to display decreased proliferation and viability (Figure 6I, J, Suppl. Figure 6F, G). We conclude that the RB family and CDKN1B/p27 are key mediators of the drug synergy between Palbociclib and Sotorasib or MRTX1133.

**Figure 6:**
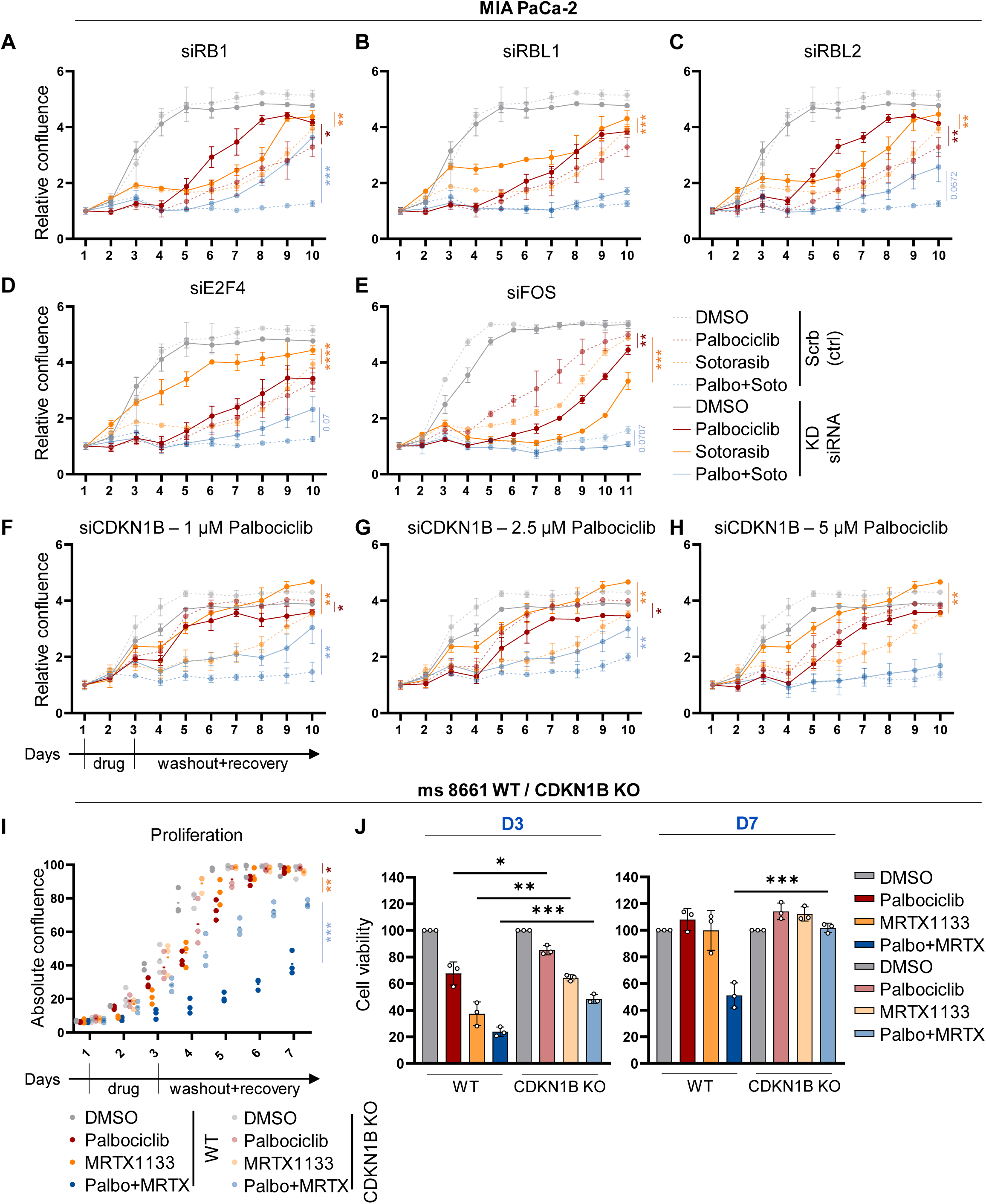
Growth suppression by inhibitors of KRAS and CDK4 depends on the RB family proteins and p27. (A) Proliferation of MIA PaCa-2 cells measured by automated transmission microscopy (Celigo®). Cells were reverse transfected by siRNAs to RB1 (A), RBL1 (B), RBL2 (C), E2F4 (D), or FOS (E); (scrb = ctrl siRNA). On day 1 after transfection, the cells were treated with DMSO, 10 µM (5 µM for (E)) Palbociclib, 5 µM Sotorasib or the combination, for 48 h, followed by seven days of recovery in normal medium. Means of three technical replicates ± SD. (F) MIA PaCa-2 cells were transfected by siRNAs to deplete CDKN1B (F, G, H) or control siRNA (scrb) during seeding. On day 1, the cells were treated with 1, 2.5 or 5 µM Palbociclib, with or without 5 µM Sotorasib, 48 h, followed by seven days of recovery without drugs. Three technical replicates, means ± SD. (I) Proliferation of 8661 cells (murine PDAC) wild type (WT) or CDKN1B knock-out (KO) lines (n=3 clones, 3 technical replicates each). All cells were treated and observed as in Fig. A, with the following specifics: 1 µM Palbociclib, 0.1 µM MRTX1133 or the combination. (J) Cell viability of 8661 WT and CDKN1B KO cells evaluated at D3 and D7 corresponding to (I). Statistical analyses: A, B, C, D, E, F, G, H, I, J unpaired t-test (A, B, C, D, E, F, G, H, I of AUC); ns: not significant, *p ≤ 0.05, **p ≤ 0.01, ***p ≤ 0.001, ****p ≤ 0.0001. Complete statistics in Suppl. Figure 6 A.

### CDK2 inhibition can substitute for antagonizing KRAS in synergy with CDK4 inhibition

Since p27 has a crucial role in mediating the drug synergy, and since p27 is predominantly inhibiting CDK2 (Polyak et al., 1994), we hypothesized that Sotorasib or MRTX1133 can be replaced by a CDK2 inhibitor when combined with the CDK4 inhibitor. Indeed, Palbociclib along with the CDK2 inhibitor Tagtociclib synergistically reduced the proliferation of MIA PaCa-2, 51T-2D, and AsPC-1 cells (Figure 7A, B, C, Suppl. Figure 7A, B, C). Furthermore, cell viability was compromised by this combination, and the Bliss synergy score confirmed the treatment to be additive to synergistic. Taken together, these observations argue that p27 acts as an endogenous cell cycle inhibitor to mediate the synergy of CDK4- and KRAS-inhibitors. Of note, however, CDK2 inhibitors cannot be expected to act specifically on tumor cells, whereas KRAS inhibitors can be tailored to act exclusively on tumor-associated KRAS mutants. Thus, in conclusion of our results, the combination of CDK4- and KRAS-inhibitors represents a promising strategy for treating KRAS-mutant cancers.

**Figure 7:**
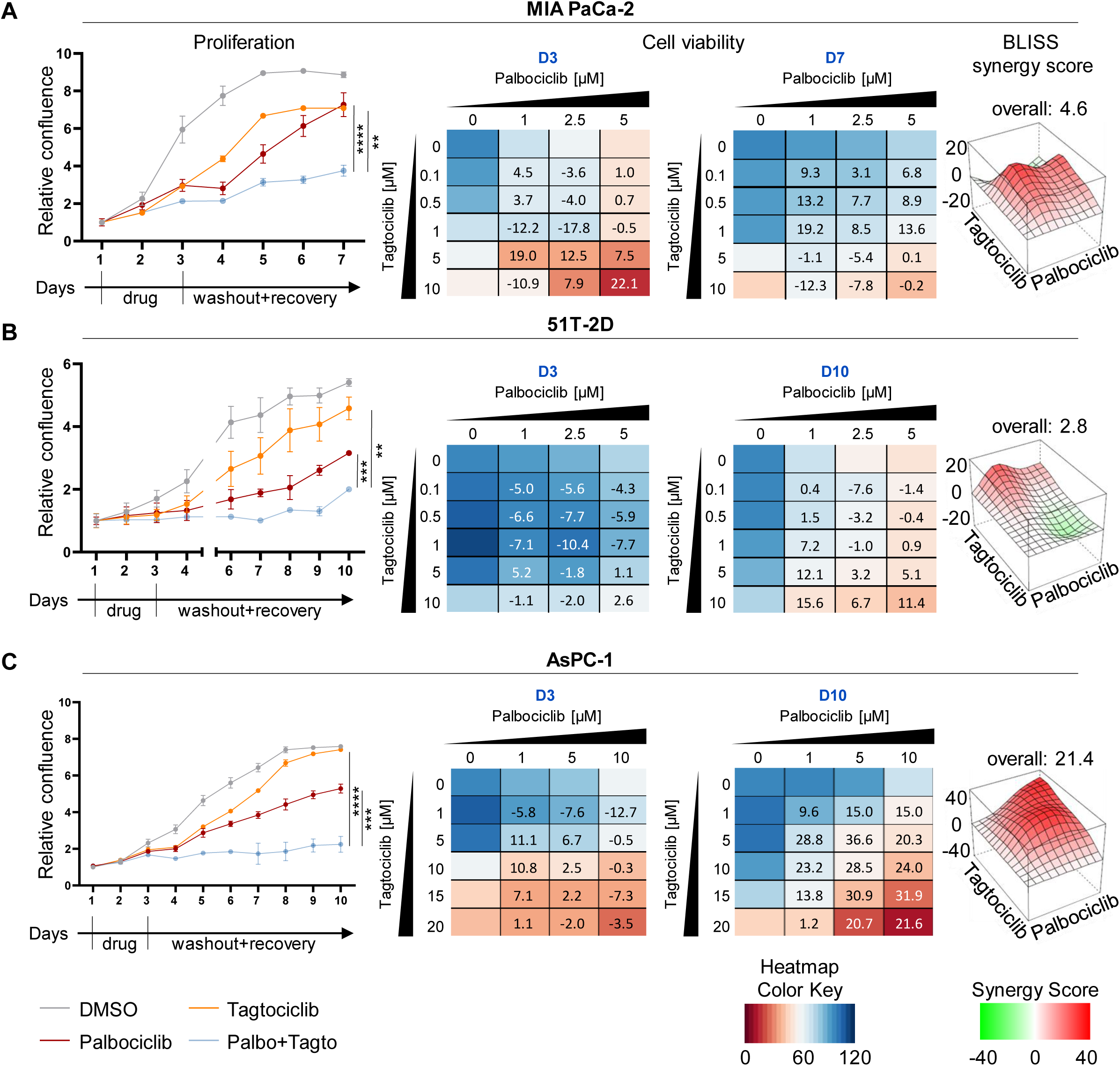
CDK2 inhibition can substitute for antagonizing KRAS in synergy with CDK4 inhibition. (A-C) All cells were treated and observed as in Fig. 1, with the following specifics: (A) Proliferation of MIA PaCa-2 cells, 5 µM Palbociclib, 1 µM Tagtociclib, evaluation at D3 and D7. (B) Proliferation of 51T-2D cells 5 µM Tagtociclib, 1 µM Palbociclib, D3 and D10. **(C)** Proliferation of AsPC-1 cells 5 µM Palbociclib, 5 µM Tagtociclib, D3 and D10.

## DISCUSSION

Our results reveal profound drug synergy against cancer cells when combining inhibitors of KRAS and CDK4. These observations were particularly striking when quantifying viable cells a few days after drug removal. Thus, the combination of the drugs confers a degree of sustainability that neither of the antagonists achieves on its own. Considering that such durable impact is particularly desirable in the context of cancer, the observations strongly argue to try these or similar combinations for therapy.

Studies performed before direct KRAS inhibitors became available to support the concept of targeting Ras signaling along with CDK4. For instance, combining inhibitors of MEK/MAP2K and CDK4 also revealed synergy (Cheng et al., 2024; Willobee et al., 2021), as did inhibitors of ERK/MAPK3 and CDK4 (Adamia et al., 2022; Goodwin et al., 2023; Jagirdar et al., 2024). Moreover, CDK4 inhibitors might improve the efficacy of drugs targeting BRAF, a kinase downstream of KRAS but upstream of MEK/ERK, e.g. in the context of malignant melanoma (Lelliott et al., 2021; Nassar et al., 2021). We have compared KRAS inhibitors with MEK inhibitors in PDAC cells (Suppl. Figure 1M, N) and observed significant but less pronounced cooperation of the latter with CDK4 inhibition. Within the limits of comparing compounds with different pharmacological specificities, this argues that direct inhibitors of KRAS act more broadly, by diminishing most if not all signaling pathways that would otherwise be driven by KRAS – besides RAF-MEK-ERK, these also include PI3K-AKT and RAL (Cox et al., 2014; Waters and Der, 2018). Perhaps, the simultaneous reduction in several pathway activities provides more opportunities for synergy with CDK4 inhibition.

Our results also suggest that RB1, through its ability to suppress E2F1 activity, is required for the efficacy of the drug combination. In addition, p27/CDKN1B acts as its mediator (Figure 8). These findings argue that tumors with RB1 deletions do not represent good candidates for treatment with CDK4 inhibitors. Mutations or deletions of *CDKN1B* are rare in cancer to begin with (Yoon et al., 2019). On the other hand, we anticipate that KRAS inhibition could be overcome by activating mutations in the downstream signaling components, e.g. the RAF kinases. However, neither RB1 nor RAF kinases are frequent subjects to mutation in PDAC (Liang et al., 2012; Witkiewicz et al., 2015), which is in support of taking the combined inhibition of KRAS and CDK4 to clinical applications for treating PDAC. However, the risk remains that potential therapies with inhibitors of KRAS and CDK4 may evoke the accumulation of such mutations in cancer cells as a result of resistance development. On top of mutations within downstream effectors, mutations within the targets themselves or within factors that act in parallel to the targets may also occur (Dobbelstein and Moll, 2014), e.g., mutations in the KRAS paralogues HRAS and NRAS, or in CDK4. On the other hand, since each of the two drugs can interfere with cell growth individually through different pathways, this may counteract the accumulation of mutations at least during the time when drug levels are sufficient.

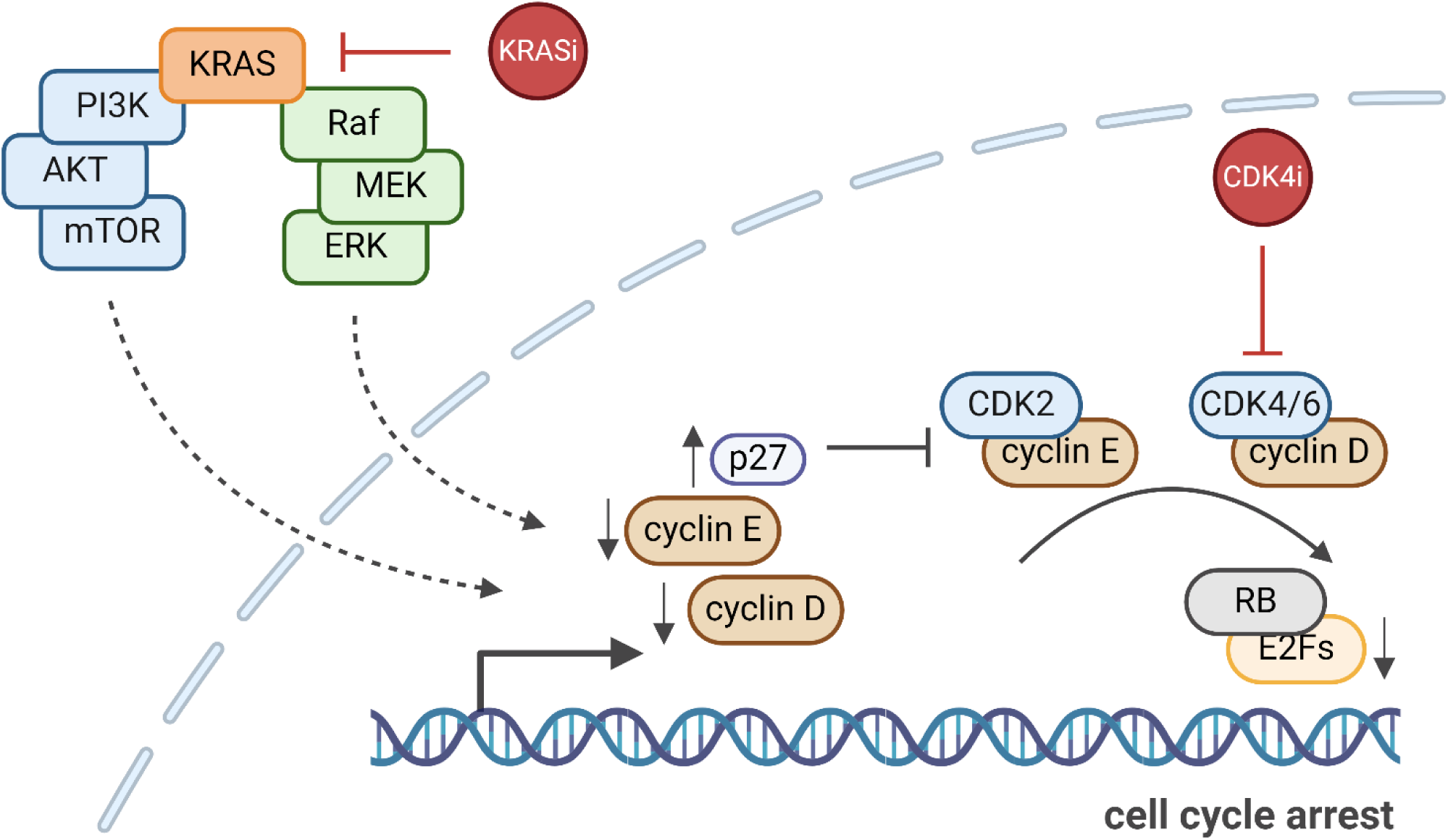

Despite the perspectives raised by this study, some limitations do apply. For instance, the induction of a senescent tumor cell phenotype may not only be beneficial to patients. Senescent cancer cells can sometimes achieve stemness and thus increased malignant potential (Milanovic et al., 2018). Moreover, the resulting inflammatory response to senescence may facilitate the proliferation of other tumor cells (Eggert et al., 2016).

With the advent of clinically applicable KRAS inhibitors, along with the long-standing establishment of CDK4 inhibitors, it has now become possible to initiate clinical applications of the findings presented here. Indeed, ongoing clinical trials are already combining inhibitors of KRAS and CDK4, albeit in early phases and along with many other combinations (NCT04185883). Potentially amenable, KRAS-mutant tumors that come to mind include not only PDACs and NSCLCs but also colorectal cancers and numerous other tumor species. This raises the perspective to set up a viable and sustainable therapeutic strategy against KRAS-mutant tumors that were hitherto notoriously difficult to treat.

## ACKNOWLEDGEMENTS

This work was supported by the German research foundation DFG, Collaborative Research Unit (CRU) 5002 (to GS, EH, MD), by the Wilhelm Sander Stiftung and the Deutsche Krebshilfe (grants to MD). M-BP was a scholar of the Göttingen graduate school GGNB, and KF received a stipend by the Else Kröner Promotionskolleg of University Medical Center Göttingen during this work. The graphical abstract was created using BioRender. We would like to thank Clara Kreutzer for her support in RNA-Isolations and protein analysis.

## AUTHOR CONTRIBUTIONS

Conceptualization: M-BP, MD

Experimentation: M-BP, KHF, CTC, CS, XF

Supervision: MD, EH, GS

Grant acquisition: MD, EH

Manuscript, initial draft: MD

Manuscript, correction and finalization: all authors

## CONFLICT OF INTEREST STATEMENT

The authors declare no conflict of interest.

## MATERIALS AND METHODS

Antibodies

| REAGENT or RESOURCE | SOURCE | IDENTIFIER |
| --- | --- | --- |
| AKT | Cell Signaling | Cat# 9272; RRID: AB_329827 |
| Alexa Flour 568 goat Anti-mouse | Thermo Fisher Scientific | Cat# A-11004; RRID: AB_2534072 |
| Alexa Fluor 488 goat Anti-rabbit | Thermo Fisher Scientific | Cat# A-11034; RRID: AB_2576217 |
| Alexa Fluor 555 goat Anti-rat | Thermo Fisher Scientific | Cat# A-21434; RRID: AB_2535855 |
| BrdU / CldU | Abcam | Cat# ab6326; RRID: AB_305426 |
| cMYC | Cell Signaling | Cat# 18583; RRID: AB_2895543 |
| Cyclin D1 | Abcam | Cat# ab134175; RRID: AB_2750906 |
| Cyclin E1 | Cell Signaling | Cat# 4129; RRID: AB_2071200 |
| Donkey anti-mouse IgG, HRP conj. | Jackson ImmunoResearch Labs | Cat# 715-036-150; RRID: AB_2340773 |
| Donkey anti-rabbit IgG, HRP conj. | Jackson ImmunoResearch Labs | Cat# 711-036-152; RRID: AB_2340590 |
| E2F-1 | Cell Signaling | Cat# 3742; RRID: AB_2096936 |
| E2F4 | Cell Signaling | Cat# 40291; RRID: AB_2799174 |
| ERK1 | Santa Cruz | Cat #sc-94; RRID: AB_2140110 |
| FOS | Proteintech | Cat#66590; RRID: AB_2881950 |
| GAPDH | Abcam | Cat# ab8245; RRID: AB_2107448 |
| HSC70 | Santa Cruz | Cat# sc-7298; RRID: AB_627761 |
| KPC2 | Abcam | Cat #ab151317 |
| KRAS | Sigma-Aldrich | Cat# WH0003845M1; RRID: AB_1842235 |
| p27 | BD Biosciences | Cat# 554069; RRID: AB_395225 |
| phospho-AKT (Ser473) (D9E) | Cell Signaling | Cat# 4060; RRID: AB_2315049 |
| phospho-ERK1/2 (Thr202/Thr204) | Cell Signaling | Cat# 4370; RRID: AB_2315112 |
| phospho-RB1 (Ser807/811) | Cell Signaling | Cat# 9308; RRID: AB_331472 |
| phospho-RB1 (T373) | Abcam | Cat# ab52975; RRID: AB_2177344 |
| RB1 (4H1) | Cell Signaling | Cat# 9309; RRID: AB_823629 |
| RBL1 | Cell Signaling | Cat# 89798; RRID: AB_2800144 |
| RBL2 | Cell Signaling | Cat# 13610; RRID: AB_2798274 |
| SKP2 | Proteintech | Cat# 15010-1-AP; RRID: AB_2187647 |
| β-actin | Abcam | Cat# ab6276; RRID: AB_2223210 |

Chemicals, peptides and recombinant proteins

| REAGENT or RESOURCE | SOURCE | IDENTIFIER |
| --- | --- | --- |
| 5-Chloro-2'-deoxyuridine | Sigma-Aldrich | Cat# C6891 |
| Abemaciclib | Selleckchem | Cat# S5716 |
| DAPI | Sigma-Aldrich | Cat# D9542 |
| DMSO | Geyer | Cat# A3672 |
| dNTP Set, 100 mM Solutions | Thermo Fisher Scientific | Cat# R0182 |
| hoechst33342 | Thermo Fisher Scientific | Cat# H3570 |
| MRTX1133 | Medchem Express | Cat# HY-134813 |
| Nuclease-Free Water | Life Technologies | Cat# AM9939 |
| Palbociclib PD 0332991 isethionate | Sigma-Aldrich | Cat #PZO199 |
| Pefabloc | Carl Roth | Cat# A154 |
| Pepstatin A | VWR | Cat# A2205 |
| Propidium Iodide | Geyer | Cat# P4864 |
| SCH772984 | Selleckchem | Cat# S7101 |
| Sotorasib | Medchem Express | Cat #HY-114277 |
| SYBR Green | Invitrogen | Cat# S7567 |
| Tagtociclib | ChemieTek | Cat# CT-PF0710 |
| Taq Polymerase | Primetech | Cat# 1800 |
| Trametinib | Selleckchem | Cat# S2673 |
| TRIzol™ reagent | Life Technologies | Cat# 15596018 |

Critical commercial assays

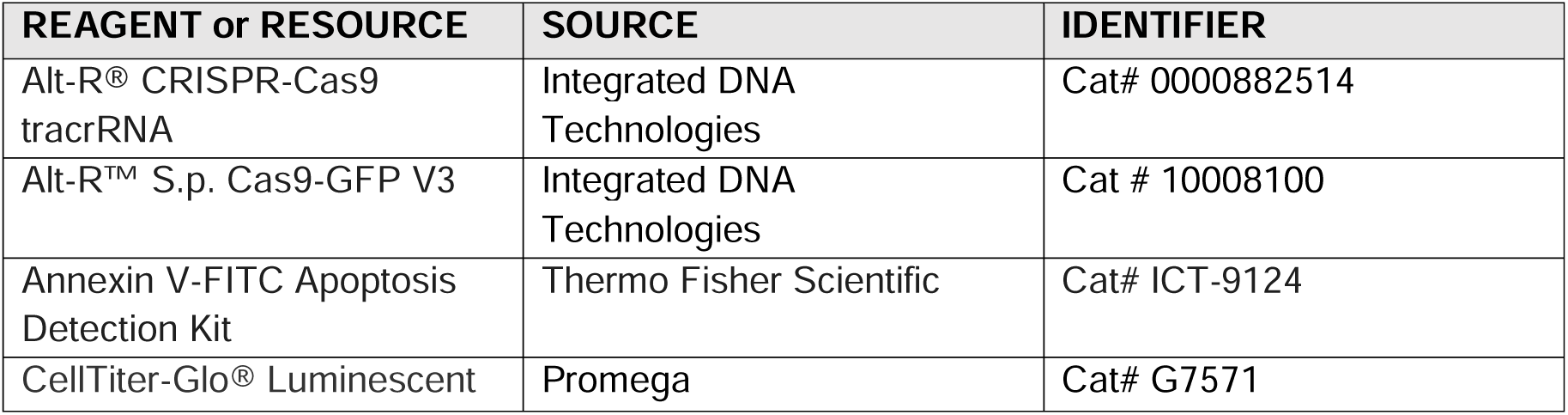
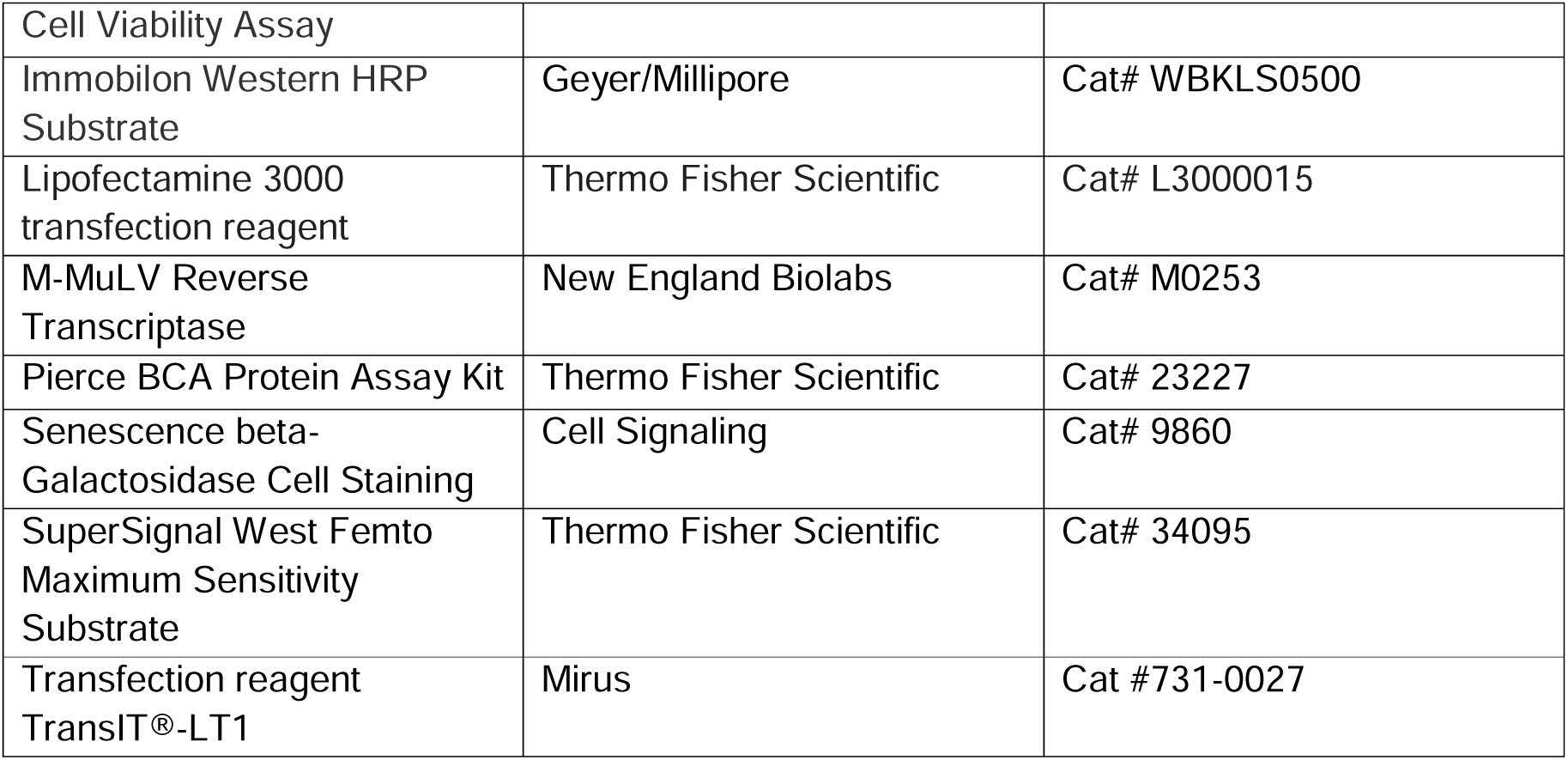

Experimental models: cell lines

| REAGENT or RESOURCE | SOURCE | IDENTIFIER |
| --- | --- | --- |
| Human: 51T-2D | Provided by Günter Schneider, CRU5002, Göttingen | N/A |
| Human: AsPC-1 | ATCC | Cat# CRL-1682; RRID: CVCL_0152 |
| Human: GöCDX52 | Provided by Elisabeth Hessman, CRU5002, Göttingen | N/A |
| Human: GöCDX53 | Provided by Elisabeth Hessman, CRU5002, Göttingen | N/A |
| Human: GöCDX7 | Provided by Elisabeth Hessman, CRU5002, Göttingen | N/A |
| Human: MIA PaCa-2 | ATCC | Cat# CRL-1420; RRID: CVCL_0428 |
| Human: NCI-H2122 | ATCC | Cat# CRL-5985; RRID: CVCL_1531 |
| Human: NCI-H358 | ATCC | Cat# CRL-5807; RRID: CVCL_1559 |
| Mouse: 8661 CDKN1B knock-out | Provided by Günter Schneider, CRU5002, Göttingen | N/A |
| Mouse: 8661 WT | Provided by Günter Schneider, CRU5002, Göttingen | N/A |
| Mouse: KPC | Provided by Elisabeth Hessman, CRU5002, Göttingen | N/A |

N/A = not available

Experimental models: organoids

N/A = not available Experimental models: organoids
| REAGENT or RESOURCE | SOURCE | IDENTIFIER |
| --- | --- | --- |
| PDO-51T (case: KFO-TM047) | Provided by CRU5002, Göttingen | N/A |

Oligonucleotides

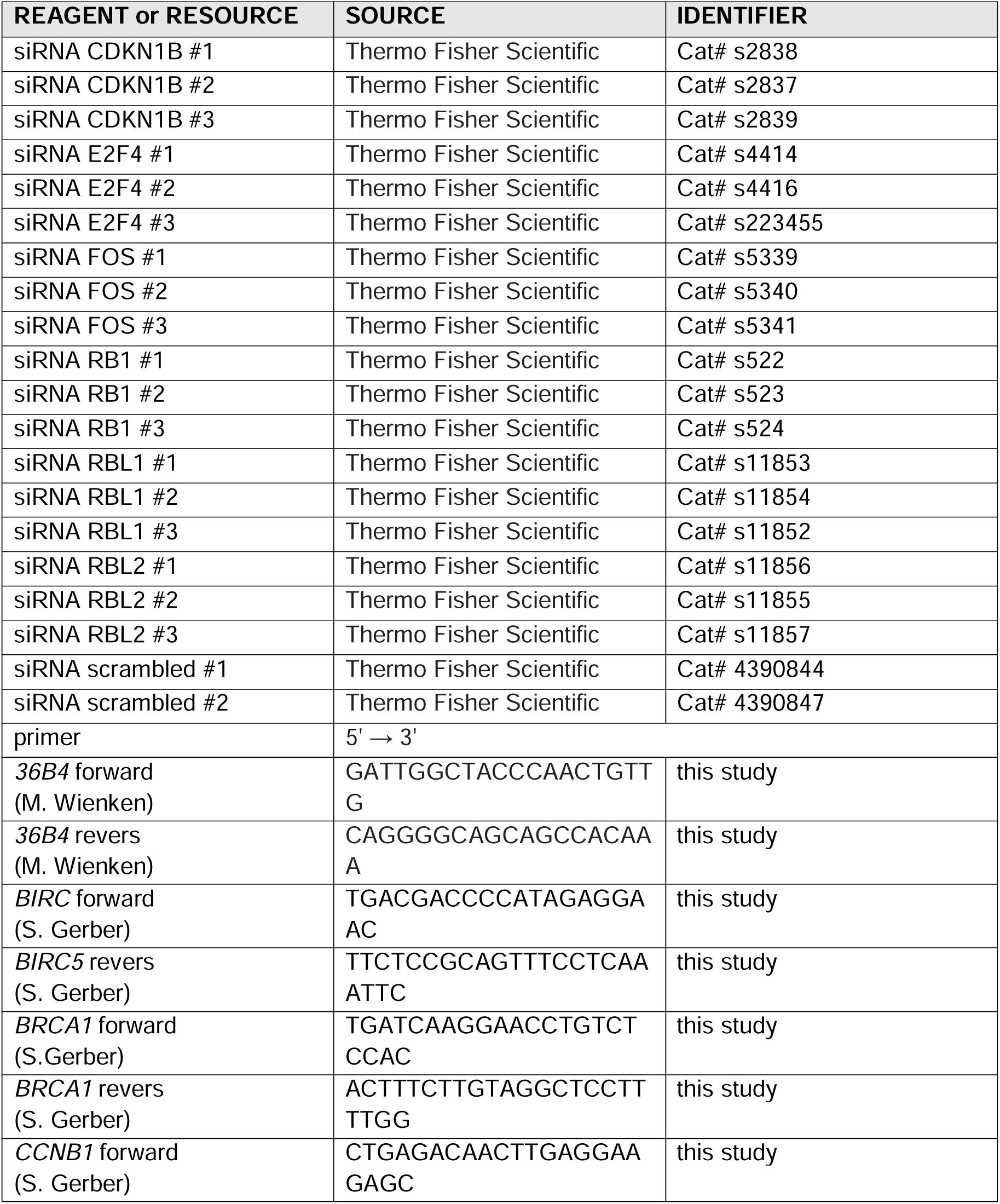
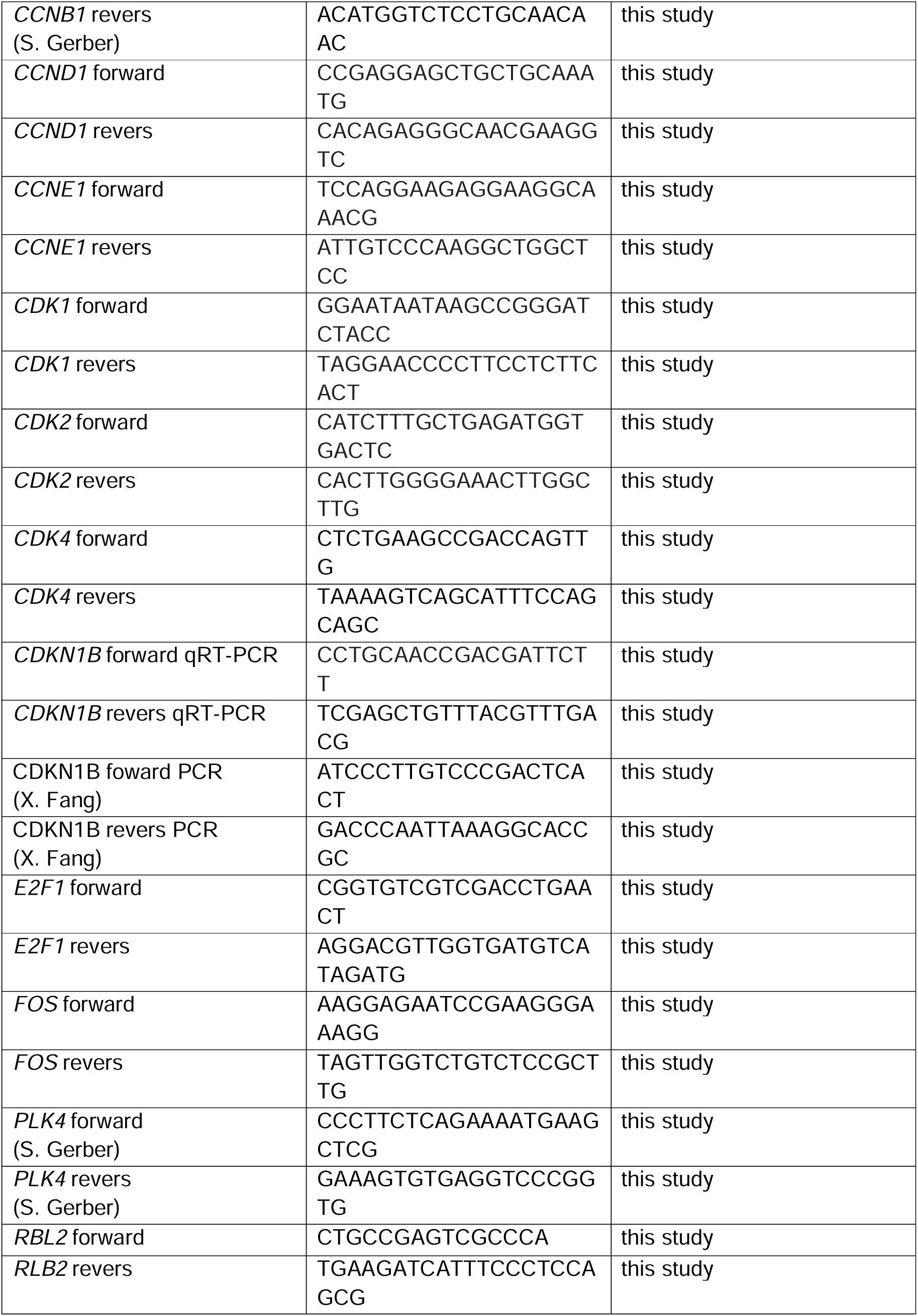

Software and algorithms

| REAGENT or RESOURCE | SOURCE | IDENTIFIER |
| --- | --- | --- |
| BioRender | BioRender | https://www.biorender.com/ |
| CeligoTM Software version2 | Nexcelom | N/A |
| FIJI version 1.54f | Image J | N/A |
| Galaxy GWDG | GWDG | http://galaxy.gwdg.de |
| Image Lab version 5.2.1 | BioRad | N/A |
| PRISM version 9+10 | GraphPad | N/A |
| R environment version 4.1.2 | www.r-project.com | N/A |
| R package: "Colorbrewer" version 1.1-2 | Neuwirt, 2022 | N/A |
| R package: "ggplots2" version 3.3.6 | Wickham et al., 2022 | N/A |
| R package: "qglots" version 3.1.3 | Warnes et al., 2022 | N/A |
| R Studio version 2022.07.02+552 | www.rstudio.com | N/A |
| Synergyfinder | synergyfinder.fimm.fi | N/A |
| ZEN Software version 3.6 | Zeiss | N/A |

### Cell culture, cell lines, and inhibitors

The human cell lines AsPC-1, NCl-H358 and HCl-H2122 were maintained in Roswell Park Memorial Institute (RPMI 1640, 42401042, life technologies) supplemented with 10% Fetal Calf Serum (FCS, ACSM0190, Anprotec), 50 U/mL Penicillin, 50 μg/mL Streptomycin (15140122, GIBCO) and 2 mM Glutamine (25030024, GIBCO). Human MIA PaCa-2 cells were maintained in Dulbecco’s modified Eagle’s medium (DMEM, 31600091, Thermo Fisher Scientific) supplemented with 10% FCS, 50 U/mL Penicillin, 50 μg/mL Streptomycin and 2 mM Glutamine. Murine KPC (LSL-KrasG12D/+; LSL-Trp53R172H/+; Pdx-1-Cre) cells were maintained in DMEM similar to MIA PaCa-2 cells with additional 1 % non-essential amino acids (11140050, Thermo Fisher Scientific). Murine 8661 cells were cultivated in DMEM GlutaMAX (61965059, GIBCO) with 10% FCS and 2 mM Glutamine. The patient-derived cell line 51T-2D was established from a patient (case: KFO-TM047) from the CRU5002 cohort (Tapia Contreras et al., 2024). 51T-2D cells were maintained in RPMI 1640 supplemented with 10% FCS and 2 mM Glutamine. Further patient-derived xenograft lines GöCDX7, GöCDX52 and GöCDX53 (cases: KFO-TM004, KFO-TM056, KFO-TM057) were maintained in a 3:1 ratio of keratinocyte-serum free medium with 25 mg bovine pituitary extract (BPE) and 2.5 µg epidermal growth factor (EGF) (17005042, GIBCO) and RPMI 1640 supplemented with 10% FCS. All cell lines were cultured at 37°C, 5% CO_2_ and regularly tested to ensure the absence of Mycoplasma contaminations. Patient-derived organoids (PDO-51T) were cultivated as described (Tapia Contreras et al., 2024).

For drug treatment, we used Abemaciclib (S5716, Selleckchem), MRTX1133 (HY-134813-5, Hycultec GmbH), Palbociclib (PZ0199-5MG, Sigma-Aldich), SCH772984 (S7101, Selleckchem), Sotorasib (HY-114277, MedChemExpress), Tagtociclib (CT-PF0710, ChemieTek) or Trametinitb (S2673, Selleckchem). DNA synthesis was quantified by 5-chloro-2-deoxyuridine (CldU) incorporation (25 µM; Sigma-Aldrich).

Transient transfections were carried out using Lipofectamine 3000 (L3000008, Thermo Fisher Scientific). siRNAs were transfected at a final concentration of 10 nM as smart pool targeting CDKN1B (s2837, s2838, s2839, Thermo Fisher Scientific), E2F4 (s4414, s4416, s223455, Thermo Fisher Scientific), FOS (s5339, s5340, s5341), RB1 (s552, s523, s524, Thermo Fischer Scientific), RBL1 (s11852, s11853, s11854, Thermo Fischer Scientific), RBL2 (s11855, s11856, s11857, Thermo Fisher Scientific); scrambled siRNA was used as control (s4390844, s4390847, Thermo Fisher Scientific).

Targeted deletion (knock-out, KO) of CDKN1B was performed in murine 8661 cells. CDKN1B was deleted by CRISPR/Cas9 using gRNA (5’-GCGGATGGACGCCAGACAAG-3’ and 5’-CAAACGTGAGAGTGTCTAAC-3’), Alt-R® CRISPR-Cas9 tracrRNA (0000882514, Integrated DNA Technologies) and Alt-R™ S.p. Cas9-GFP V3 (10008100, Integrated DNA Technologies) through lipofection (TransIT®-LT1, Mirus, Lot No. 12094394). 48 h after lipofection, the media was changed, and green fluorescent protein (GFP) signals were checked through the microscope. 24 h after the media change, single-cell cloning was conducted by seeding WT and CDKN1B KO cells into 96-well plates (0.75 cells per well). 15 h after seeding, the wells were manually checked for the existence of single cells in each well. Wells with more than one cell were excluded from further handling. After 2-3 weeks of cultivation, clones were expanded and analysed. To select clones containing the desired deletion, dot western blot was performed to exclude clones with p27 protein expression. In addition, PCR was performed using the forward (5’-ATCCCTTGTCCCGACTCACT-3′) and reverse primers (5’-GACCCAATTAAAGGCACCGC-3′). The targeting of CDKN1B was confirmed by Sanger sequencing of the PCR product.

### Cell viability assays

The Cell Titer-Glo® Luminescent Cell Viability Assay (G7571, Promega) was used to determine the viability of adherent cells. Viability was plotted using R environment and R Studio (www.r-project.com, www.r-studio.com, version 4.2.1) with the following packages “Colorbrewer” ((Neuwirth, 2022) version 1.1-2), “ggplots2” ((Wickham et al., 2022) version 3.3.6) and “gplots” ((Warnes et al., 2022) version 3.1.3). The synergy of two drugs was calculated via synergyfinder.fimm.fi (Ianevski et al., 2017) using the BLISS score and interpreted as follows. Bliss score >10: synergy; -10 – 10: additive mechanism; <-10: antagonism.

### Cell proliferation assays

Adherent cells were seeded into 96well plates (3606, Corning) and treated with drugs the following day for 48 h. Then, the cells were incubated with their respective media. Cell confluence was measured every 24 h by bright field microscopy using the Celigo^TM^ Cytometer (Nexcelom), for up to 11 days. Confluence was determined with the corresponding Celigo^TM^ software (Nexcelom, software version 2.0) by analysing the percentage of surface occupied by adherent cells over the total surface.

### Immunoblot analyses

Whole cell lysates were prepared in RIPA lysis buffer, i.e. 20 mM Tris-HCl pH 7.5, 150 mM NaCl, 10 mM EDTA, 1% Triton-X 100, 1% sodium deoxycholate, 0.1% SDS, 2□M urea, and protease inhibitors (pepstatin, leupeptin hemisulfate, aprotinin, AppliChem) and sonicated for 10 min. The BCA protein assay kit (23227, Thermo Fisher Scientific) was used to determine the total protein concentration. Samples were boiled for 5 min at 95°C in Laemmli buffer, and equal amounts of proteins were separated by SDS-PAGE, followed by transfer to a nitrocellulose membrane. The membrane was blocked with 5 % (w/v) non-fat milk (Roth) in TBS supplemented with 0.1% Tween-20 (AppliChem) for 2 h. Primary antibody incubation was performed at 4°C overnight. Peroxidase (HRP)-conjugated secondary antibodies (Jackson ImmunoResearch Labs) were applied to the membranes, and proteins were detected using Immobilon Western Substrate (WBKLS0500, Millipore) or Super Signal West Femto Maximum Sensitivity Substrate (34095, Thermo Fisher Scientific).

### Reverse transcription and real-time quantitative PCR (qRT-PCR)

Total RNA was extracted from cells using TRIzol® (15596018, Life Technologies). 200 µL Chloroform was added per 1000 µL TRIzol. For phase separation, samples were centrifuged, and the aqueous phase was used to precipitate the RNA with 800 µL isopropanol. Two washing steps with 75 % ethanol were performed before the RNA was resuspended in nuclease-free water (AM9939, Life Technologies). The concentration was determined via spectrophotometry.

RNA was reverse-transcribed using oligo-dT and random nonamers as primers, followed by qRT-PCR analysis using SYBR Green (S7567, Invitrogen), as previously described (Jansen et al., 2024) . Gene expression levels were normalized to the mRNA levels from the housekeeping gene *36B4*, and the analysis was conducted using the ΔΔCt method.

### Transcriptome analyses

Total mRNA of 51T-2D and AsPC-1 cells was obtained as described above. Sequencing of mRNA (≥ 400 ng) was performed on Illumina NovaSeq 6000 by Novogene (read length: paired-end 150 bp). mRNA sequencing data was processed in the Galaxy environment provided by the GWDG (Gesellschaft für wissenschafltiche Datenverarbeitung mbH Göttingen, https://galaxy.gwdg.de/). In short, a quality check was performed using FastQC (version 0.69). Afterwards, the first 11 nucleotides of each read were trimmed (Trim, version 0.0.2). Fastq files were aligned to the hg19 reference genome using HISAT2 (version 2.0.5.2). Read counts for each sample and gene were acquired via featureCounts (version 1.6.3+galaxy2). Finally, DeSeq2 (version 2.11.40.6+galaxy1) was used to perform differential gene expression analysis and to obtain normalized counts (Suppl. Table 1, 2). To analyse differentially regulated genes, only genes with a cut-off of |log2fold| ≥ 0.6, adjusted p-value (padj.) < 0.05, and baseMean ≥ 15 were included. Gene Set Enrichment Analysis of hallmark gene sets (GSEA, version 4.1.0, https://www.gsea-msigdb.org/gsea/msigdb/index.jsp) was used to identify impaired pathways. Differentially expressed genes were correlated with the Molecular Signature Database (MsigDB) using the Enrichr platform (https://maayanlab.cloud/Enrichr/). Z-Score heat maps of differentially regulated genes were obtained from Morpheus (https://software.broadinstitute.org/morpheus/).

### Nucleoside analogue incorporation assay

Cells were seeded in 96well plates (3606, Corning) and treated as indicated. 25 µM CldU was applied to cells for 2 h at 37°C, 5 % CO_2_ before fixation with 4 % PFA (Sigma-Aldrich) in PBS for 20 min. Cells were washed 4x in PBS. Permeabilization and denaturation was performed using 0.5 % Triton-X-100/2.5 M HCl for 1 h. Blocking was carried out with 3 % BSA (w/v) + 0.1 % Tween-20 in PBS for 1 h. This was followed by incubation with a primary antibody (α-BrdU, ab6326, Abcam) over night at 4°C. After three washing steps with blocking solution, the cells were incubated with the secondary antibody (A-21434, Thermo Fisher Scientific), together with 4′,6-diamidino-2-phenylindole (DAPI, D9542, Sigma-Aldrich) for 1 h at RT. Images were acquired using the Cell Discoverer 7® (Zeiss) with 20x magnification. ZEN software (Zeiss, version 3.6) was used to quantify the CldU signal over the DAPI signal.

### Immunofluorescence staining of organoids

Organoids were seeded in 96 wells plates (3606, Corning) and treated as indicated. On day 7, the organoids were incubated with Hoechst 33342 (100 μg/mL, H3570, Thermo Fisher Scientific) and Propidium Iodide (PI, P4864, Geyer). Images were acquired using the Cell Discoverer 7® (Zeiss) with 20x magnification. Brightfield and immunofluorencence images were acquired as Z-stacks to depict complete organoids.

### Annexin V assay

Cell death was analysed by performing Annexin V staining. The Annexin V-FITC Apoptosis Detection Kit (ICT-9124, Biozol) was used, with the addition of PI (10 μg/mL, P4864, Geyer), Hoechst 33342 (100 μg/mL, H3570, Thermo Fisher Scientific). To quantify cell death, cells were seeded in low density and stained at the indicated time points. Analysis was performed using the Celigo^TM^ Cytometer (Nexcelom) with the corresponding Celigo^TM^ software (Nexcelom, software version 2.0).

### Cell senescence determination by senescence-associated beta-galactosidase (SAB) staining

Cell senescence was analysed using the Senescence beta-Galactosidase Cell Staining kit (9860, Cell Signaling). First, the cells were washed and fixed. The staining solution was prepared freshly for every use and was applied over night at 37 °C. Imaging was performed using a Zeiss Axiocam 503 color microscope. Finally, FIJI was used to analyse ten images per condition with the script imagej_cell_count_jython (https://github.com/lquenti/imagej_cell_count_jython).

### Statistics

Statistical testing was performed using GraphPad Prism Software (Version 10). Unpaired t-tests were used for statistical analysis of area-under-curve (AUC, cell proliferation analysis), ordinary one-way ANOVA followed by Turkey’s multiple comparison was used for RT-PCRs, immunofluorescence staining, and organoid viability analysis. Ns, not significant; ∗p-value≤0.05; ∗∗p-value≤0.01; ∗∗∗p-value≤0.001; ∗∗∗∗p-value□<□0.0001. All statistical details can be found in the figure legends.

---

**Suppl. Figure 1:**
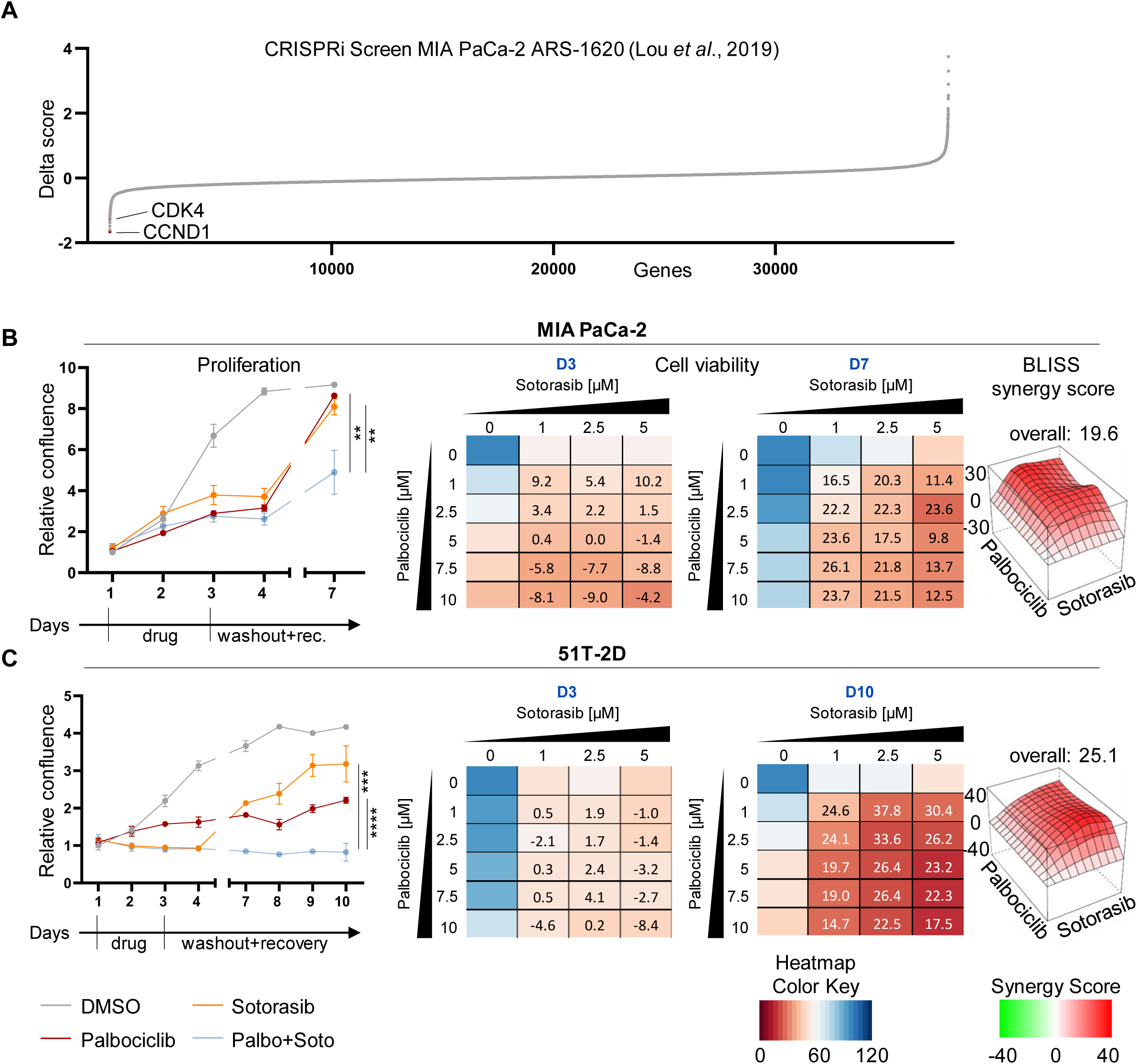
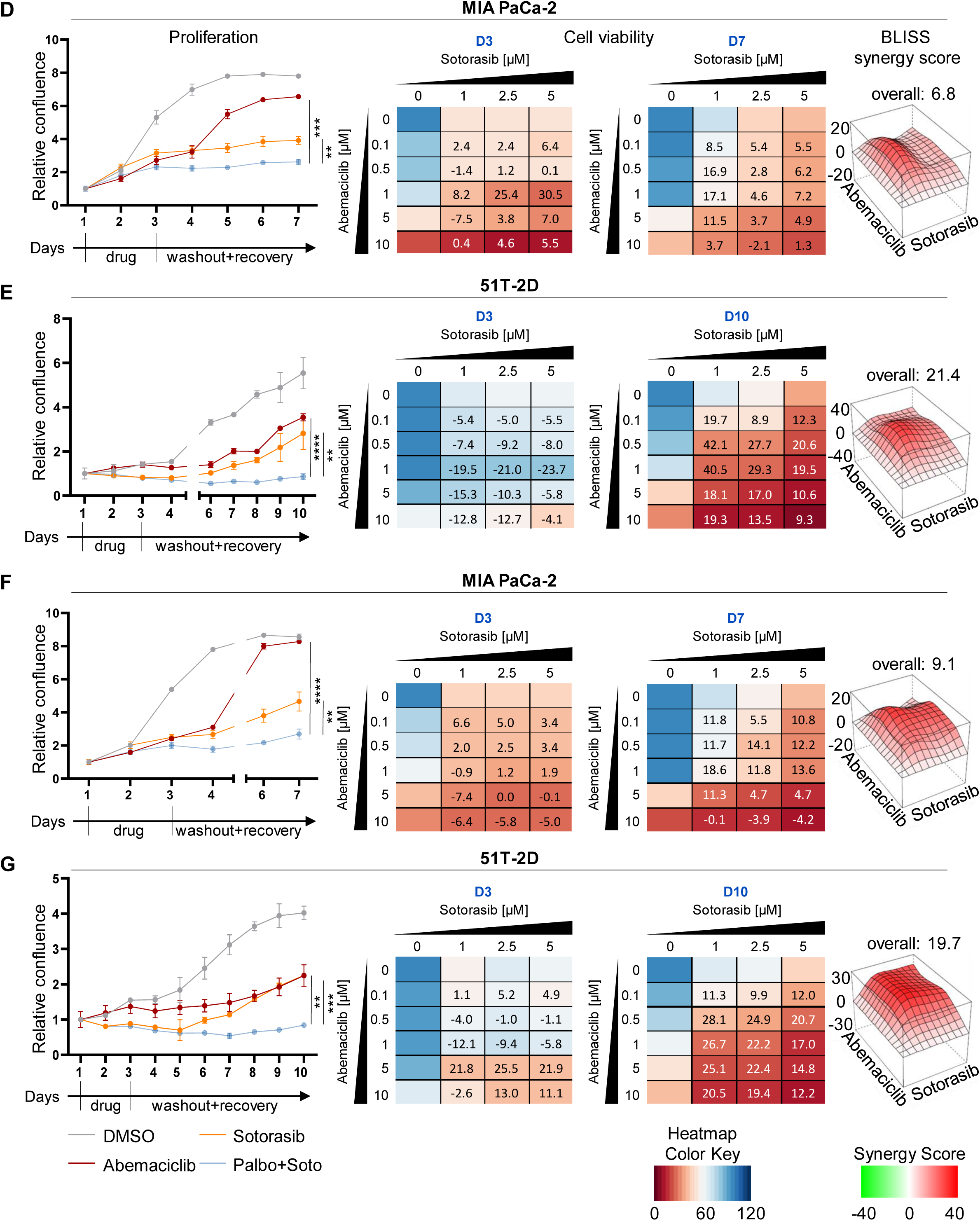
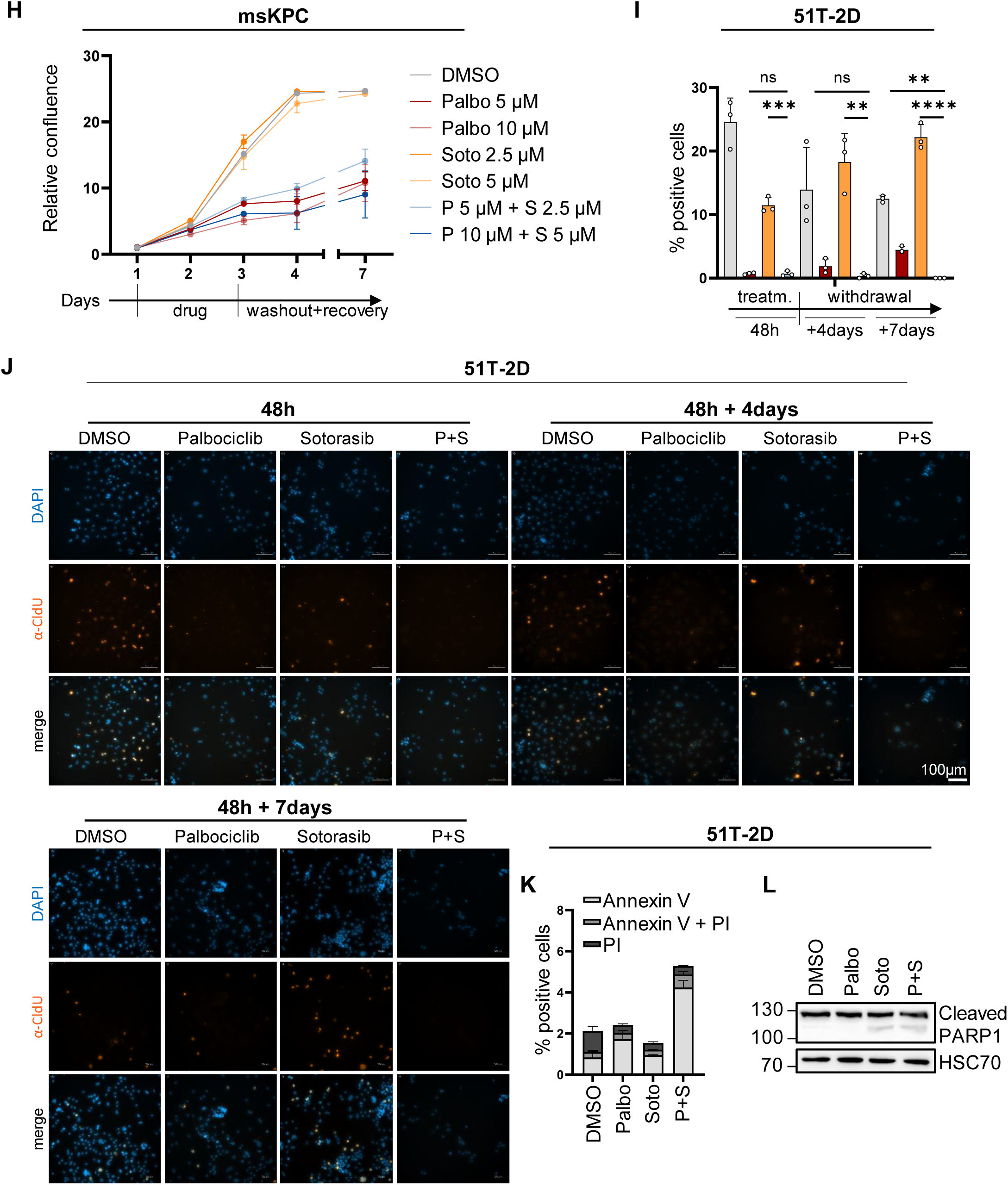
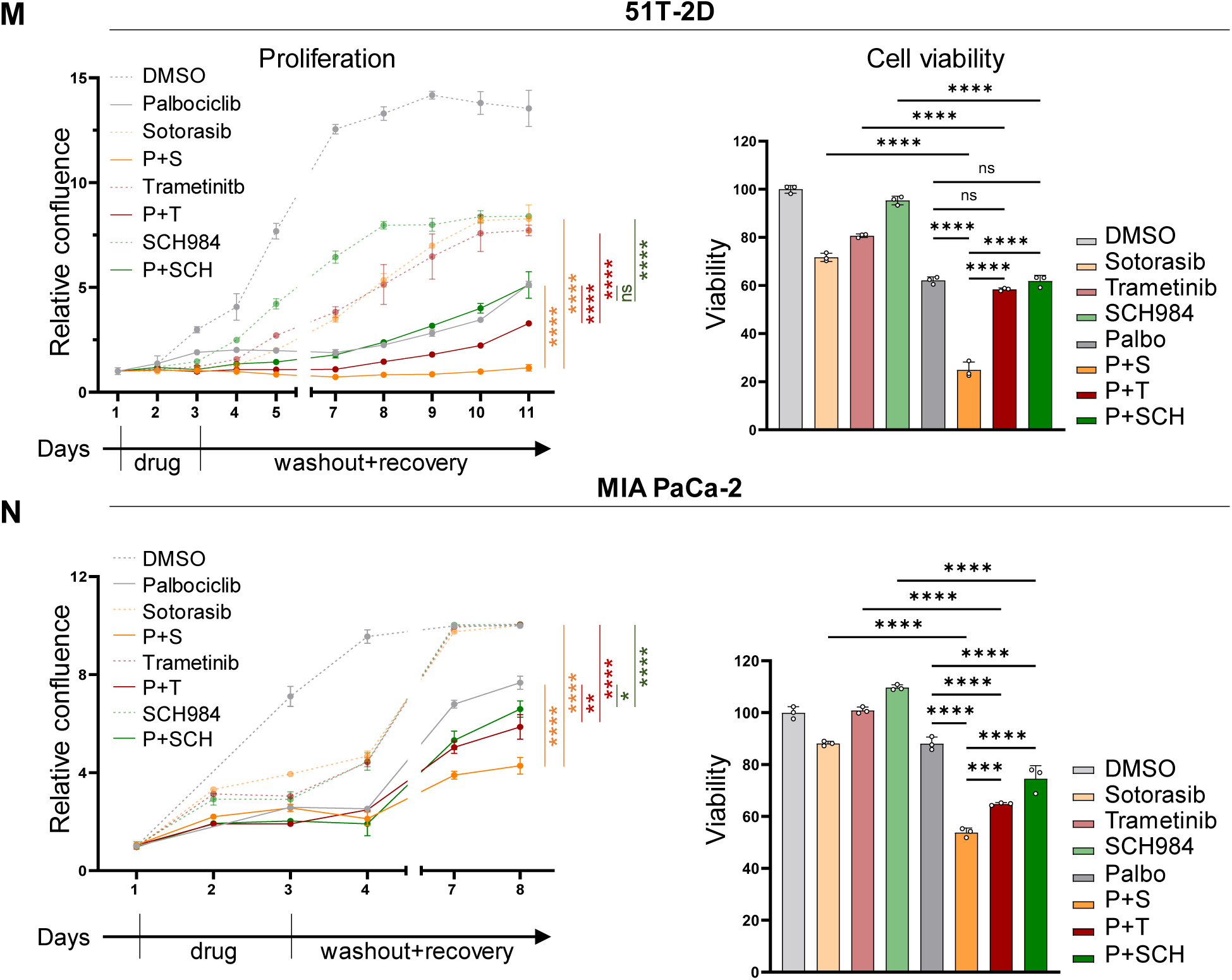
The KRAS mutant G12C-targeting drug Sotorasib synergizes with Palbociclib to suppress the growth of pancreatic cancer cells and organoids, with sustainability after drug removal. (A) CRISPRi screen of MIA PaCa-2 cells treated with 300 nM ARS-1620 in comparison to DMSO (Lou *et al*., 2019). Analysed Delta score of Suppl. Table 4. CDK4 and CCND are highlighted in red. (B) (left) Proliferation of MIA PaCa-2 cells, measured by automated microscopy (Celigo®) upon treatment with DMSO, 5 µM Palbociclib and 2.5 µM Sotorasib or the combination for 48 h, followed by four days of recovery in normal medium. Graphs indicate the mean of three technical replicates ± SD. (right) Heat maps depict the viability of MIA PaCa-2 cells right after treatment (D3) and after recovery (D7), normalized to the DMSO control. Bliss synergy scores were determined based on cell viability at D10. (C) 51T-2D cell proliferation was determined as in (B); 5 µM Palbociclib and 2.5 µM Sotorasib; D3 and D10 BLISS synergy map at D10. (D) MIA PaCa-2 cells, 1 µM Abemaciclib and 5 µM Sotorasib, D3 and D7 BLISS synergy map at D7. (E) 51T-2D cells, 0.5 µM Abemaciclib and 1 µM Sotorasib, D3 and D10. (F) MIA PaCa-2 cells, 1 µM Abemaciclib and 5 µM Sotorasib, D3 and D7. (G) 51T-2D cells, 0.5 µM Abemaciclib and 1 µM Sotorasib, D3 and D10. (H) KPC cells, 5/10 µM Palbociclib and/or 2.5/5 µM Sotorasib for 48 h, followed by four days of recovery in normal medium. Means of three replicates ± SD. (I) 51T-2D cells treated with 5 µM Palbociclib, 2.5 µM Sotorasib or both, for 48 h, with subsequent washout and further incubation for four or seven days. The cells were stained with DAPI and α-CldU to indicate DNA synthesis. CldU positive cells were quantified using the Celldiscoverer7®. (J) Representative images to (I) are shown. Scale bar 100 µm. (K) Apoptosis quantification using the Celigo®. 51T-2D cells were as in (I) for 48 h followed by seven days of recovery in normal medium. For analysis of apoptosis cells were stained with Annexin V and PI. Quantification was carried out via the Celigo®. (L) Immunoblot analysis of whole-cell lysates of 51T-2D. Cells were treated as in (I) for 48 h. HSC70 served as loading control. (M) 51T-2D cells, 5 µM Palbociclib, 2.5 µM Sotorasib, 25 nM Trametinib or 0.5 µM SCH772984. Bar graphs show cell viability right after treatment (D3) and after recovery (D11), normalized to the DMSO control. (N) MIA PaCa-2 cells, 10 µM Palbociclib, 2.5 µM Sotorasib, 25 nM Trametinib or 0.5 µM. Cell viability as in (M) at D3 and D11. Statistical analyses: B, C, D, E, F, G unpaired t-test; I, M, N one-way ANOVA followed by Tukey’s multiple comparison (B, C, D, E, F, G, M, N: of AUC); ns: not significant, *p ≤ 0.05, **p ≤ 0.01, ***p ≤ 0.001, ****p ≤ 0.0001.

**Suppl. Figure 2:**
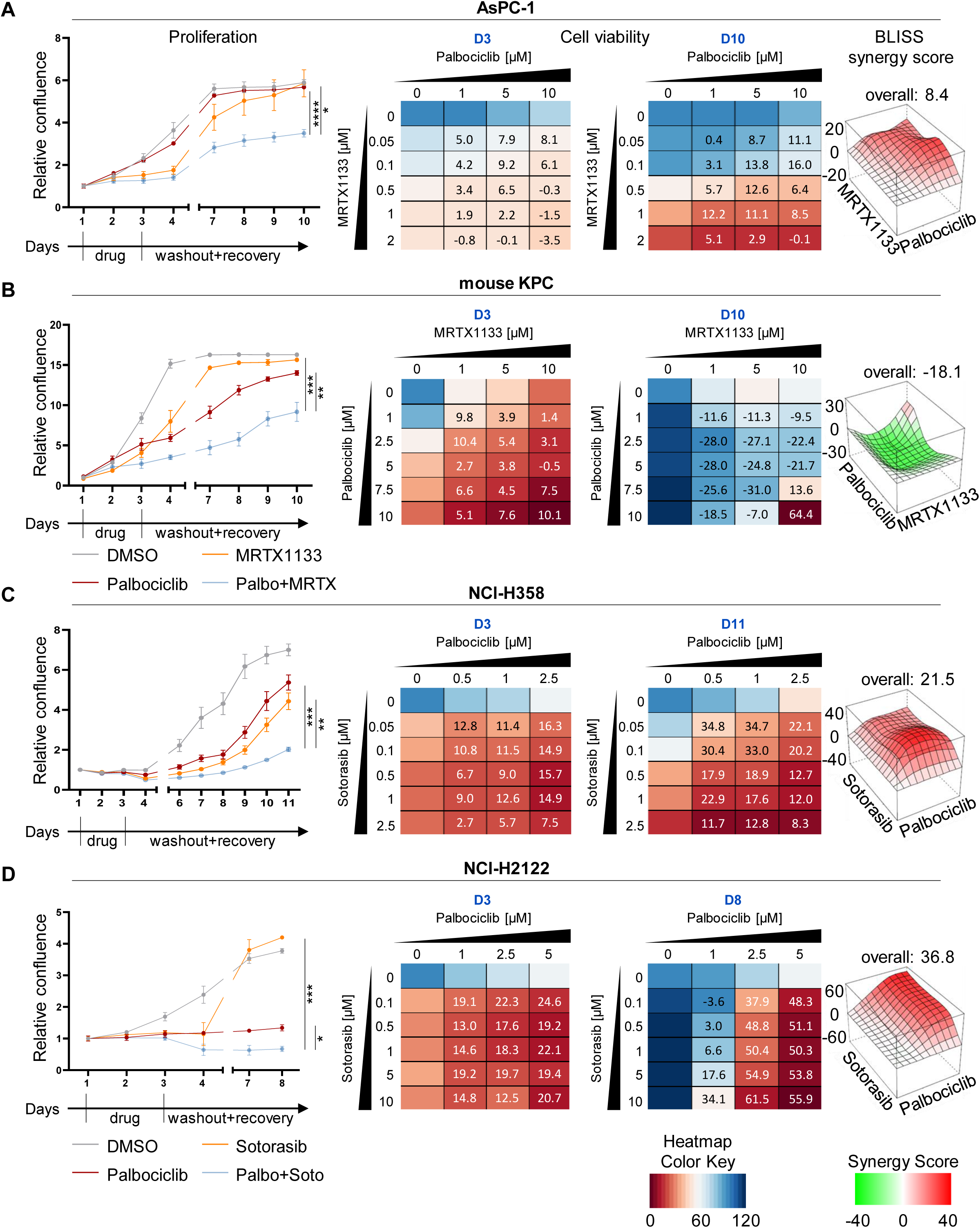
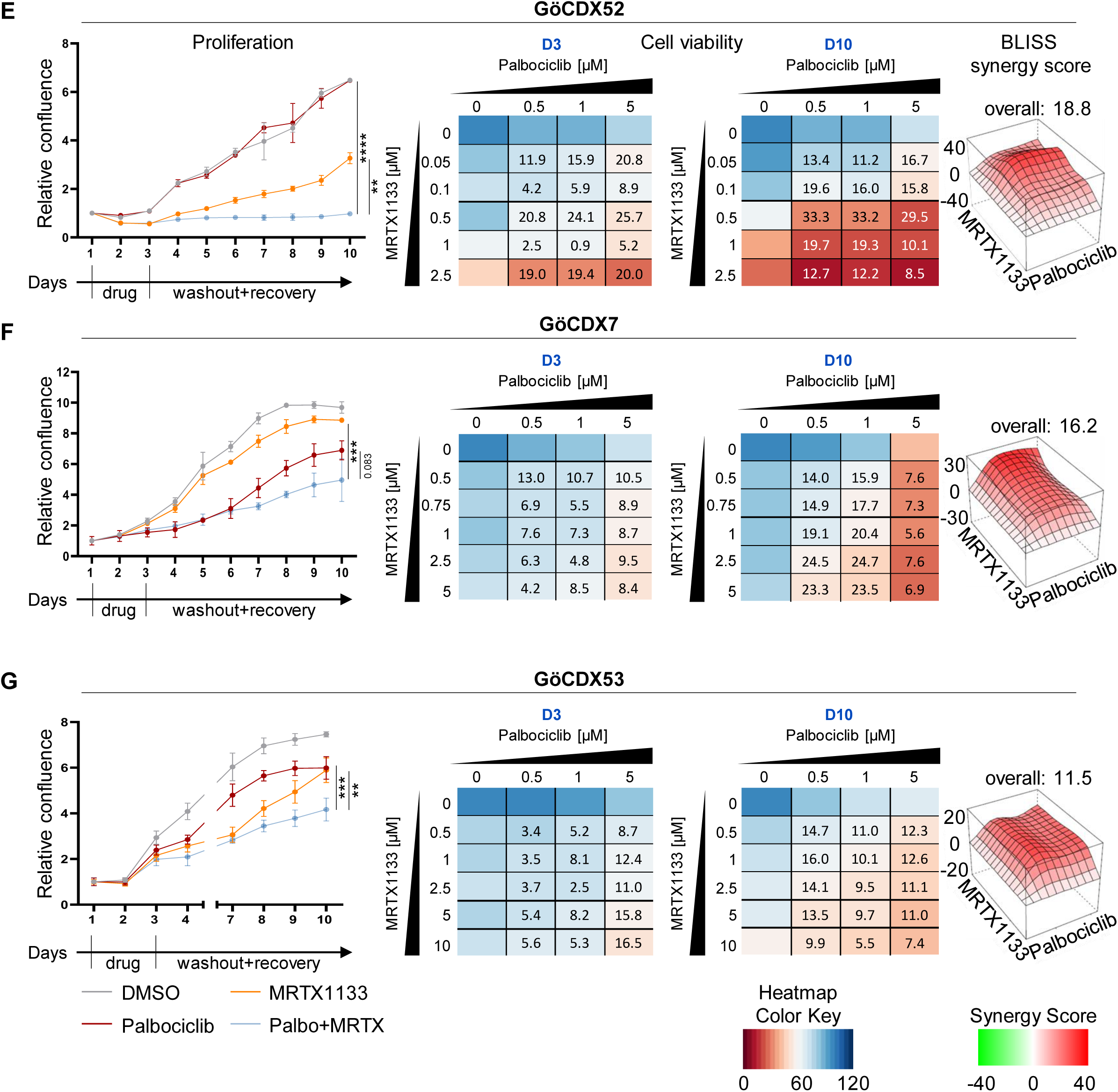
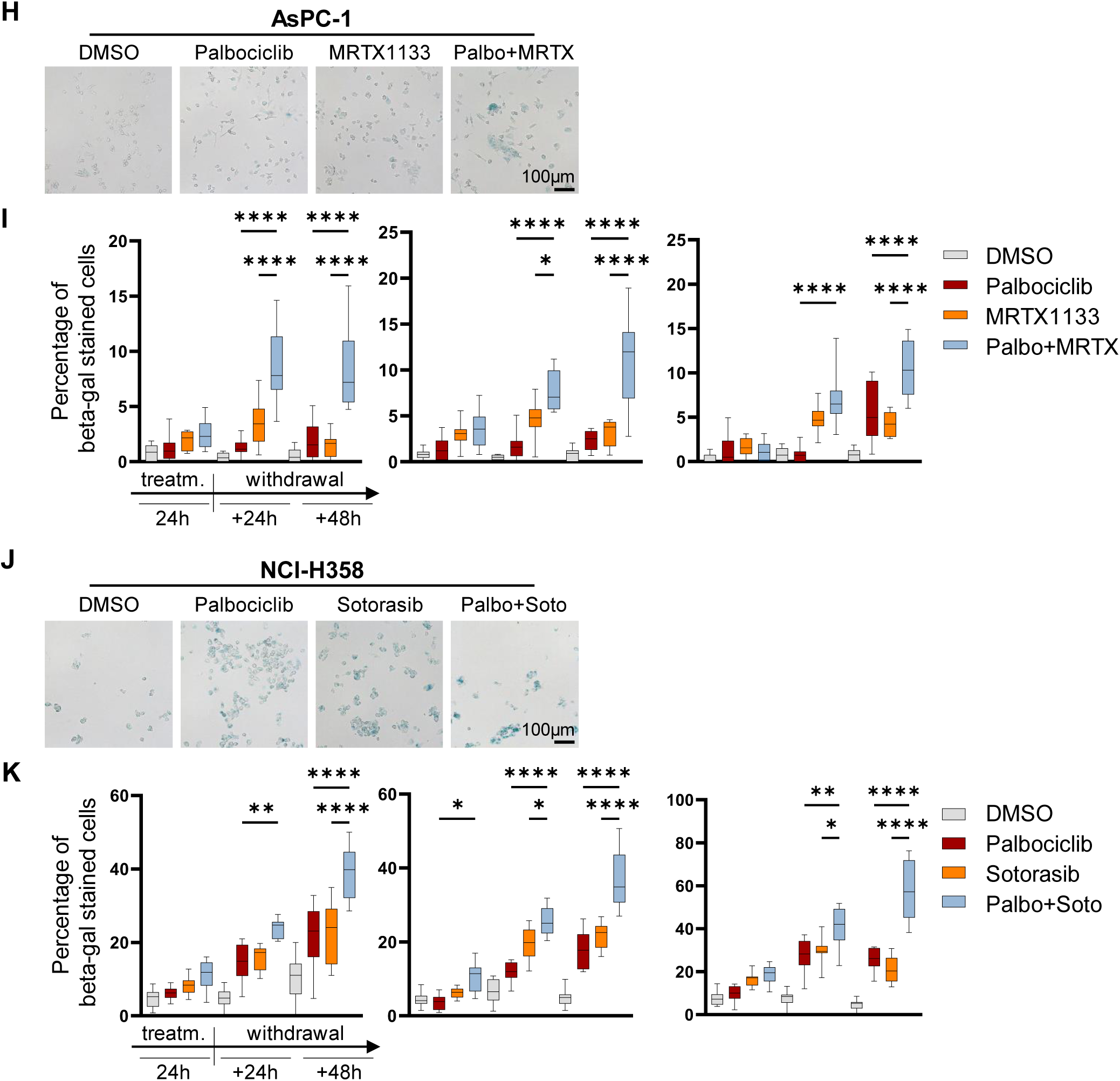
Palbociclib and KRAS inhibitors synergize to suppress PDAC and NSCLC cell proliferation, and to induce senescence. All assays were carried out as described in the legend to Suppl. Fig. 1. (A) AsPC-1 cells, 5 µM Palbociclib and 0.1 µM MRTX1133, D3 and D10. BLISS scores were determined for D10. (B) KPC cells, 5 µM Palbociclib and 5 µM MRTX1133, D3 and D10. (C) NCl-H358 cells, 0.5 µM Palbociclib and 50 nM Sotorasib, D3 and D11. (D) NCl-H2122 cells, 5 µM Palbociclib and 10 µM Sotorasib, D3 and D8. (E) GöCDX52 cells, 1 µM Palbociclib, 0.5 µM MRTX1133, D3 and D10. (F) GöCDX7 cells, 0.5 µM Palbociclib, 0.5 µM MRTX1133, D3 and D10. (G) GöCDX53 cells, 5 µM Palbociclib, 5 µM MRTX1133, D3 and D10. (H) Representative images of senescence associated beta-galactosidase (SAB) staining in AsPC-1 cells treated with DMSO, 5 µM Palbociclib, 0.1 µM MRTX1133 or the combination for 24 h, followed by 48 h of recovery in normal medium. Brightfield images: sharpen: 75 %, brightness -25 %, contrast +50 %, saturation 120 %. Scale bar 100 µm. (I) Quantification of beta-galactosidase positive cells was performed using ImageJ. 10 Images were quantified and are shown as mean ± SD; one representative replicate is shown, n=3. (J) SAB staining in NCl-H358 cells treated with DMSO, 1 µM Palbociclib, 0.1 µM Sotorasib or the combination for 24 h, followed by 48 h without drugs. Image settings as in (H). (K) Quantification of NCI-H358 cell senescence as in (I), n=3. Statistical analyses: A, B, C, D, E, F, G unpaired t-test; I, K one-way ANOVA followed by Tukey’s multiple comparison (A, B, C, D, E, F, G: of AUC); ns: not significant, *p ≤ 0.05, **p ≤ 0.01, ***p ≤ 0.001, ****p ≤ 0.0001.

**Suppl. Figure 3:**
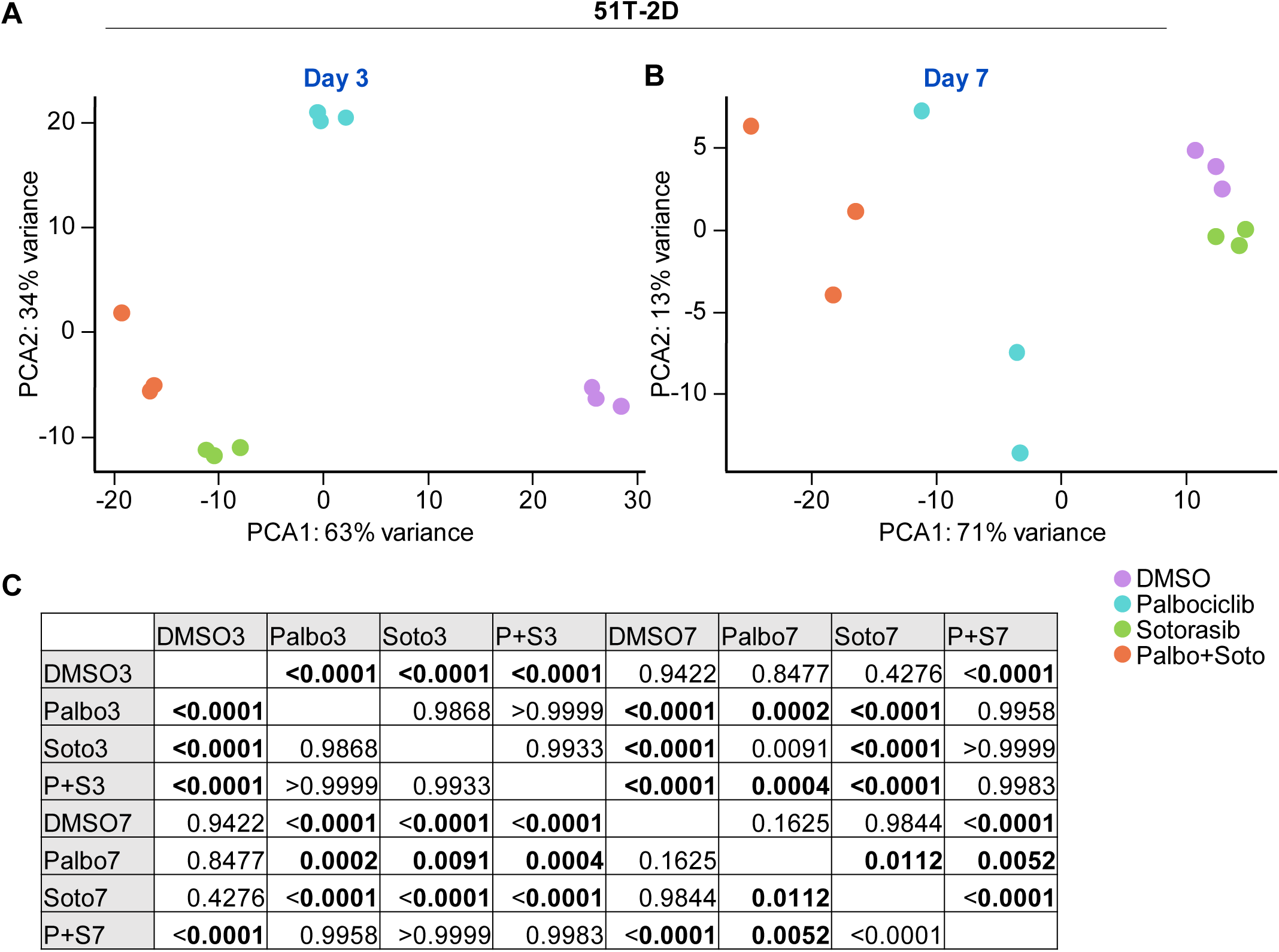
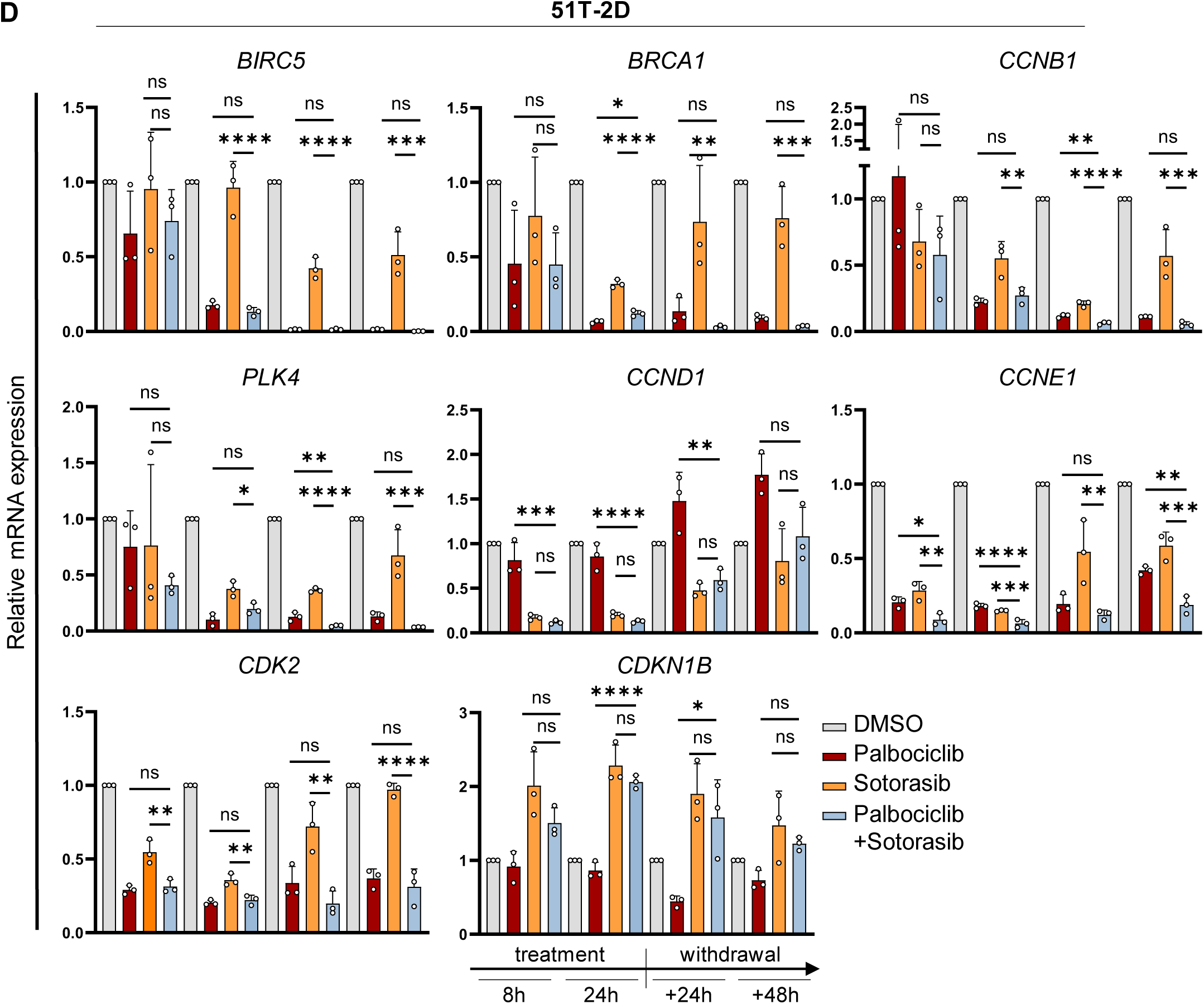
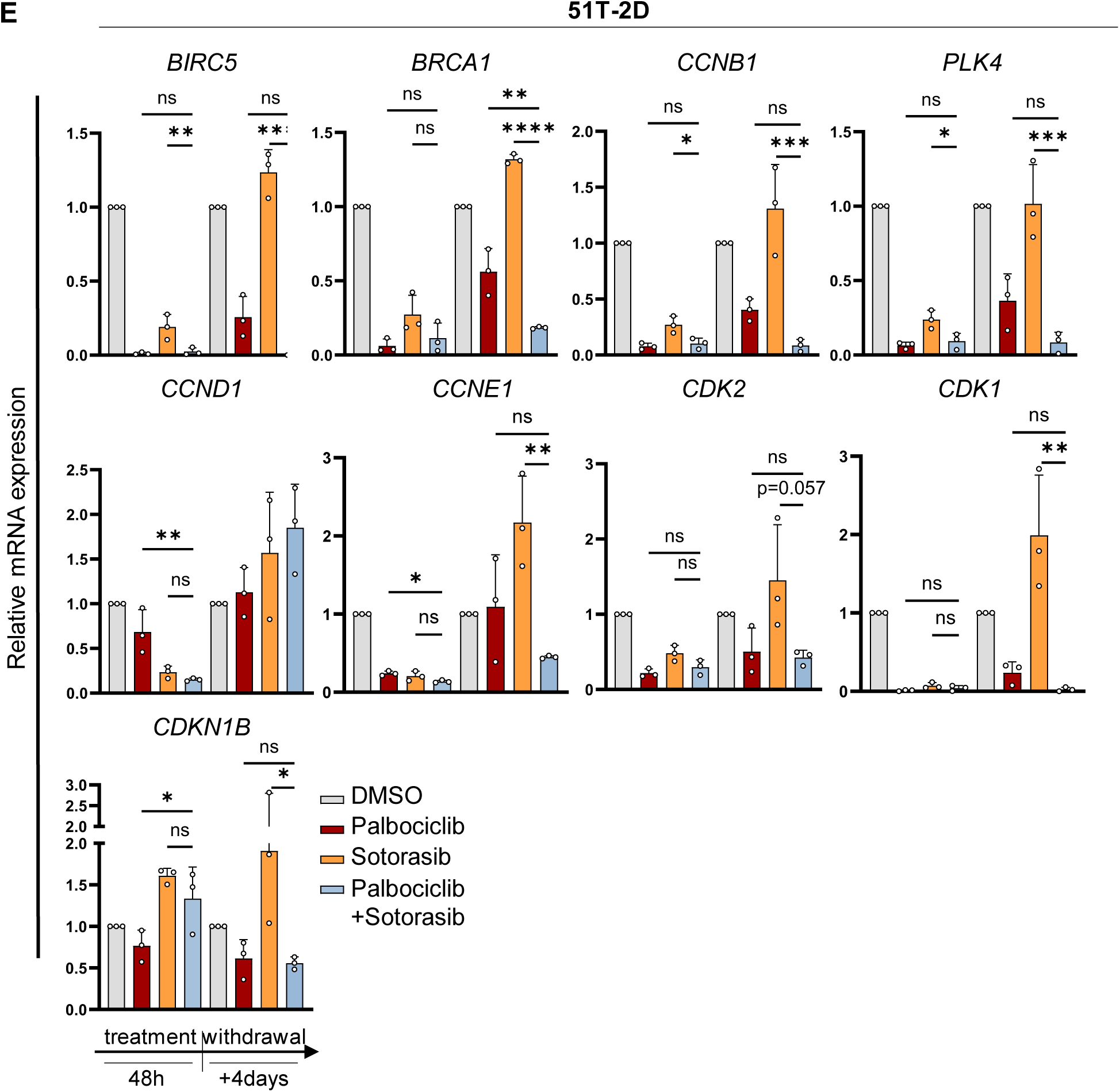
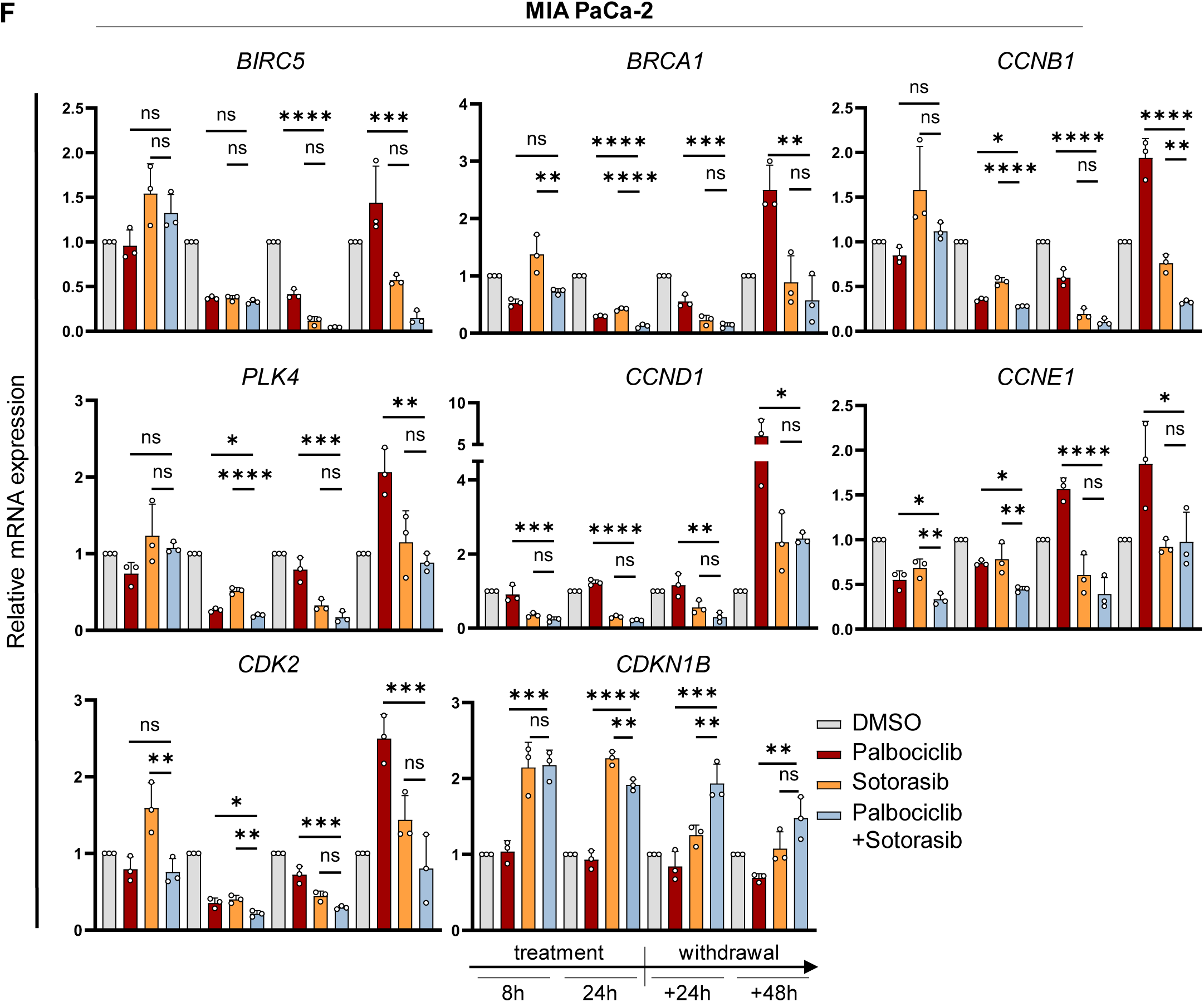
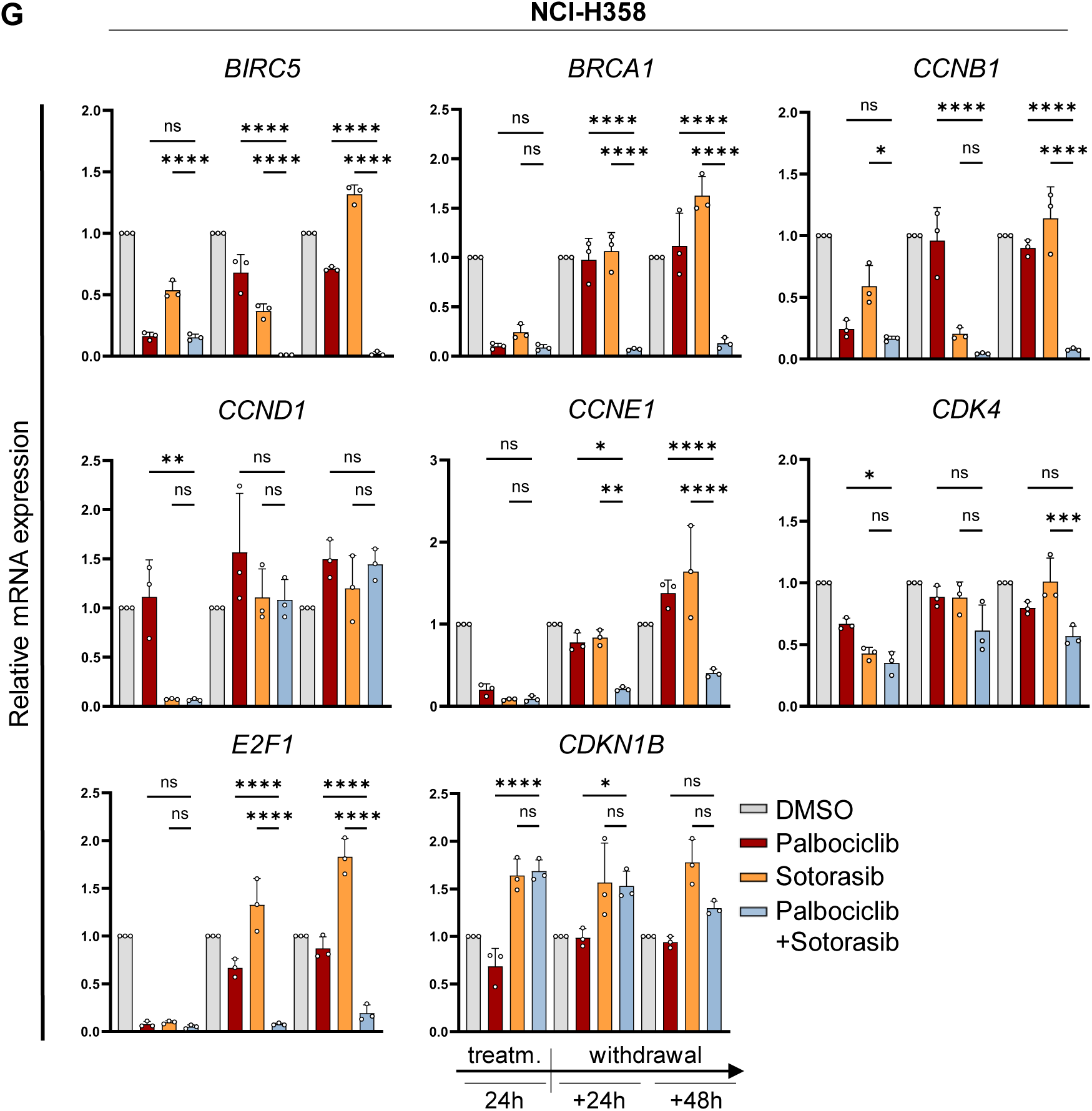
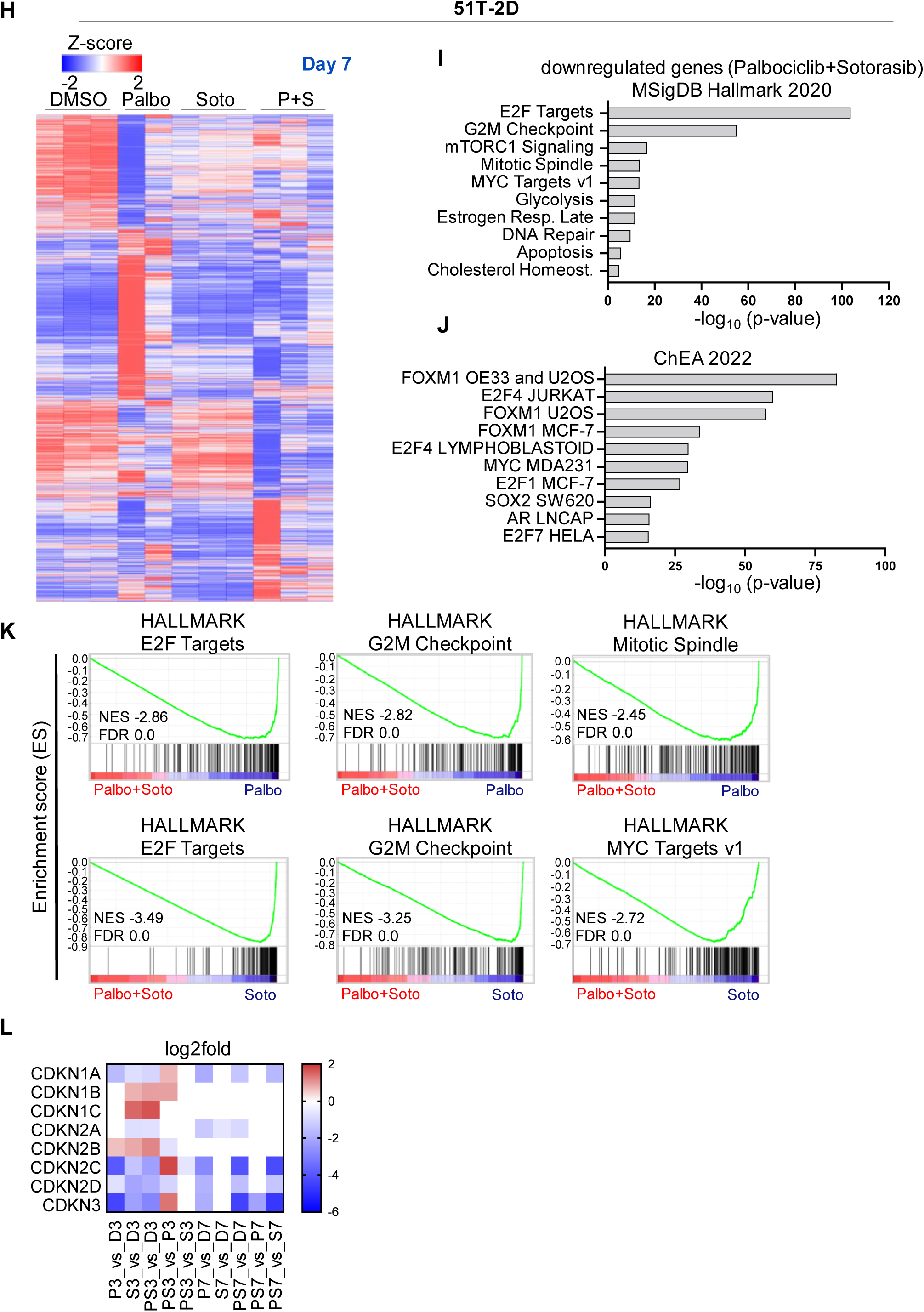
Inhibitors of KRAS G12C and CDK4 cooperate to switch the transcriptome towards ceased proliferation. (A) PCA plot of RNA-seq data derived from 51T-2D cells revealed clustering of treatment on day 3 (48 h of treatment) and (B) day 7 (48 h of treatment + 4 days washout). Upper replicate of Palbociclib day 7 was excluded (Palbociclib day 7: n=2), rest n=3. (C) Statistical analyses: one-way ANOVA followed by Tukey’s multiple comparison of Fig. 3E. (D) Expression of E2F target genes and cell cycle regulators in 51T-2D cells treated with DMSO, 5 µM Palbociclib, 2.5 µM Sotorasib or the combination of both for 8 and 24 h and washout for 24 or 48 h. Expression levels were normalized to *36B4* mRNA and shown as mean ± SD. (E) Expression of genes in 51T-2D cells treated and analysed as in (D) for 48 h and washout for four days. (F) Expression of genes in MIA PaCa-2 cells treated with 10 µM Palbociclib, 5 µM Sotorasib or the combination of both for 8 and 24 h and washout for 24 or 48 h, determined as in (D). (G) Expression of genes in NCl-H358 cells treated with 1 µM Palbociclib, 0.1 µM Sotorasib or the combination of both for 24 h and washout for 24 or 48 h, determined as in (D). (H) Heat map depicting differentially expressed genes according to the z-score after performing DeSeq2 analysis four different samples (DMSO, 5 µM Palbociclib, 2.5 µM Sotorasib or combination treatment for 48 h + 4 days washout (Day7), n=3) of 51T-2D cells. Only genes with |log2fold| ≥ 0.6, adjusted p-value (padj.) < 0.05, and baseMean ≥ 15 were included in the analysis. (I) Downregulated genes upon treatment with Palbociclib + Sotorasib (48 h treatment + 4 days washout) vs DMSO were correlated with the Molecular Signature Database (MSigDB) Hallmark 2020 and ChEA 2022 (J) dataset using the Enrichr platform to identify potentially impaired pathways. (|log2fold| ≥ 0.6, padj. < 0.05, baseMean ≥ 15) Top 10 (ChEA 2022: human), p-value ranked (-log10). (K) Gene set enrichment analysis (GSEA) of combination treatment vs Palbociclib or Sotorasib after 48 h of treatment + 4 days washout (D7) of Hallmarks (h.all.v2023.2). (L) Expression of cyclin dependent kinase inhibitors after 48 h treatment (3) or 48 h treatment + 4 days of drug withdrawal (7): |log2fold| ≥ 0.6. Statistical analyses: D, E, F, G one-way ANOVA followed by Tukey’s multiple comparison; ns: not significant, *p ≤ 0.05, **p ≤ 0.01, ***p ≤ 0.001, ****p ≤ 0.0001.

**Suppl. Figure 4:**
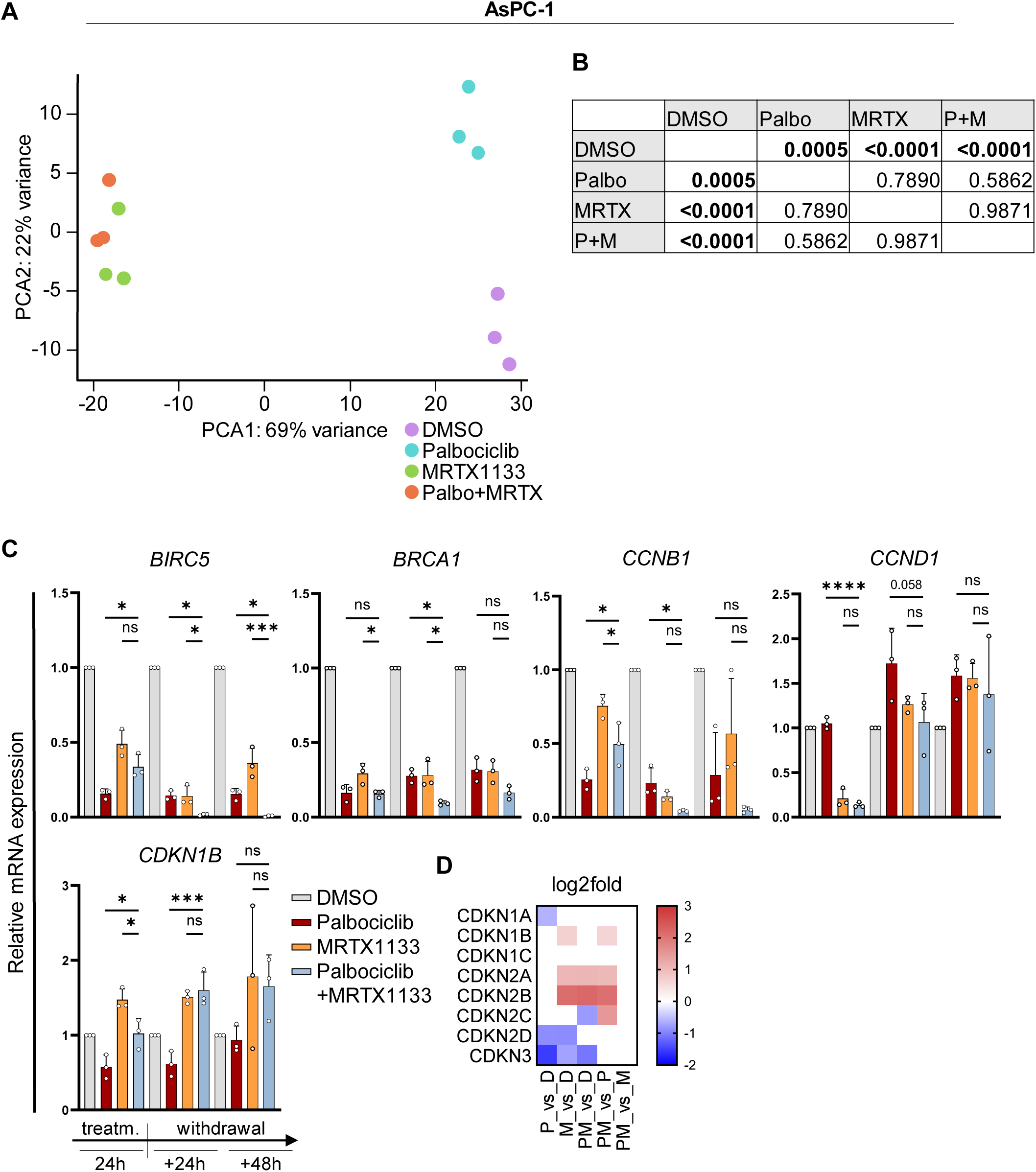
Transcriptome alterations in PDAC cells treated with Palbociclib and MRTX1133.) (A) PCA plot of RNA-seq data derived from AsPC-1 cells indicating clustering of the treatment groups on day 2 (24 h of treatment), n=3. (B) Statistical analyses: one-way ANOVA followed by Tukey’s multiple comparison of Fig. 4E. (C) Expression of E2F target genes and cell cycle regulators in AsPC-1 cells treated with 5 µM Palbociclib, 0.5 µM MRTX1133 or the combination of both for 24 h and washout for 24 or 48 h. Expression levels were normalized to *36B4* mRNA and shown as mean ± SD. (D) Expression of cyclin dependent kinase inhibitors after 24 h treatment: |log2fold| ≥ 0.6. Statistical analyses: B, C one-way ANOVA followed by Tukey’s multiple comparison; ns: not significant, *p ≤ 0.05, **p ≤ 0.01, ***p ≤ 0.001, ****p ≤ 0.0001.

**Suppl. Figure 5:**
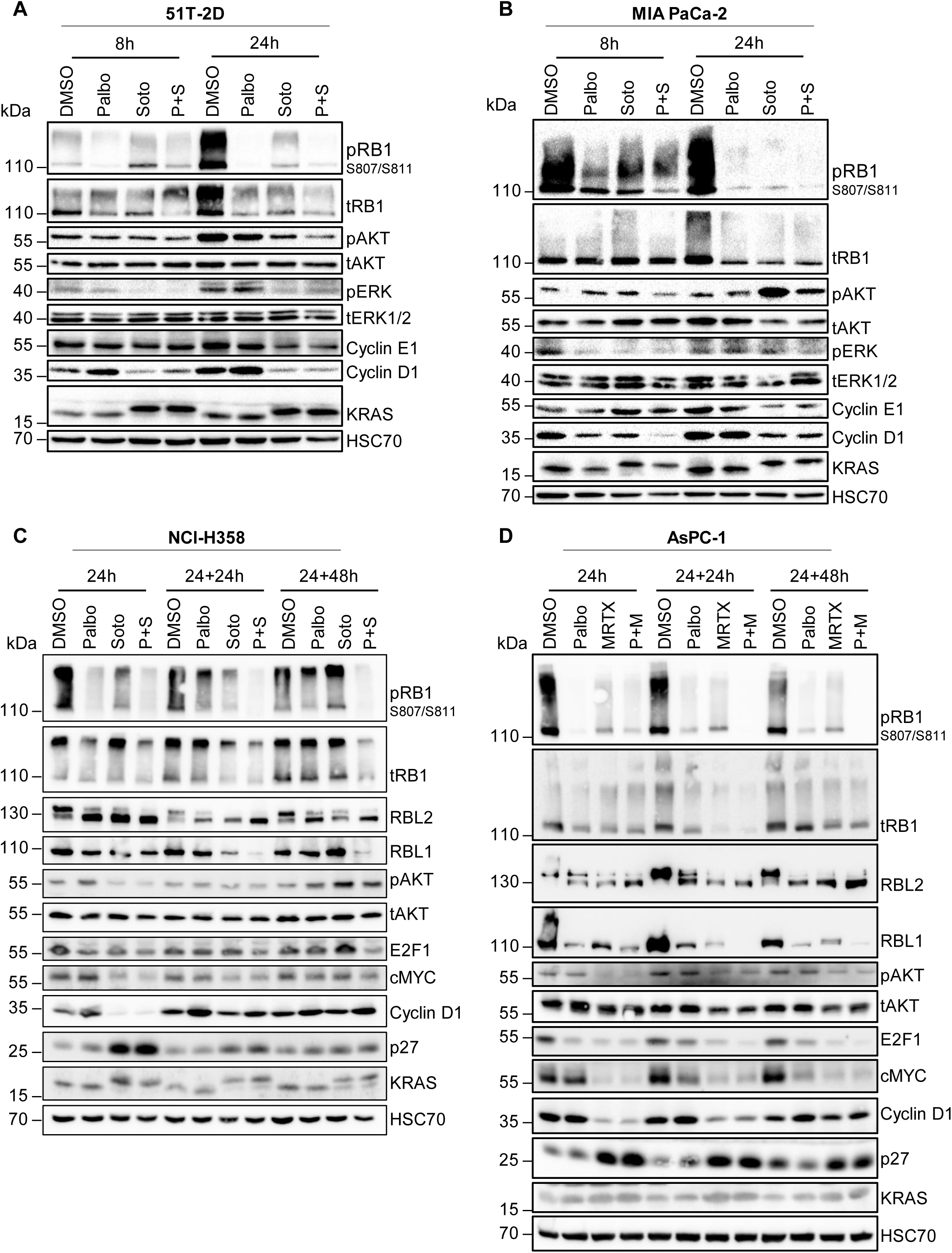
Upon inhibition of KRAS and CDK4, E2F1 largely disappears while CDKN1B/p27 levels increase and RB family proteins adopt a hypophosphorylated state. (A) Immunoblot analysis of whole-cell lysates of 51T-2D cells treated with DMSO, 5 µM Palbociclib, 2.5 µM Sotorasib or combination treatment for 8 or 24 h. HSC70 served as sample control. One representative immunoblot shown; n=2. (B) MIA PaCa-2 cells treated with 10 µM Palbociclib and/or 5 µM Sotorasib for 8 or 24 h. n=2. (C) NCl-H358 cells treated with 1 µM Palbociclib and/or 0.1 µM Sotorasib for 24 h and washout for 24 or 48 h. n=2. (D) AsPC-1 cells treated with 5 µM Palbociclib and/or 0.5 µM MRTX1133 for 24 h and washout for 24 or 48 h. n=2.

**Suppl. Figure 6:**
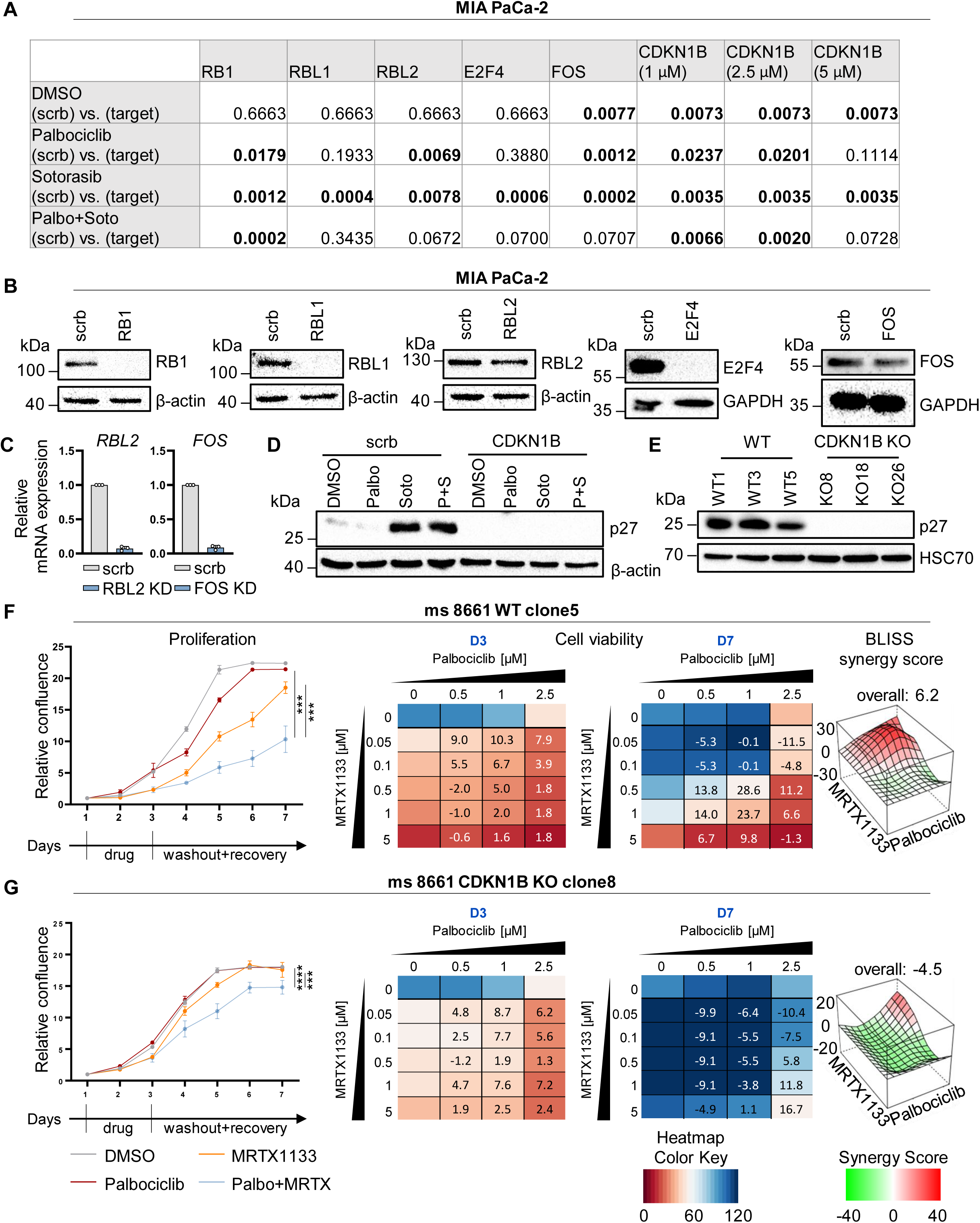
Growth suppression by inhibitors of KRAS and CDK4 depends on the RB family proteins and p27. (A) Statistical analyses corresponding to Fig. 6 A, B, C, D, E, F, G, H unpaired t-test of. Bold: p < 0.05; scrb = control siRNA; target = target siRNA. (B) Immunoblot analysis of MIA PaCa-2 cells corresponding to Fig. 6 A, B, C, D, E confirming knockdowns. HSC70/GAPDH/β-actin served as loading control. (C) Expression of *RBL2/FOS* in MIA PaCa-2 cells after knockdown of *RBL2 or FOS* corresponding to Fig.6 C, E. Scrb siRNA transfected cells served as controls. Data was normalized to *36B4* mRNA and is shown as mean ± SD, n=3. (D) Immunoblot analysis of MIA PaCa-2 cells corresponding to (Fig. 6 F, G, H) confirming knockdowns. β-actin served as loading control. (E) Immunoblot analysis of 8661 cells CDKN1B WT clones 1, 3, 5 and CDKN1B knock-out KO clones 8, 18, 26 confirming the knock-out of CDKN1B. (F) 8661 WT (F) and CDKN1B knock-out (G) were treated and observed as in Suppl. Fig. 1, with the following conditions: 1 µM Palbociclib, 0.5 µM MRTX1133 or the combination. Evaluation at D3 and D7. Statistical analyses: A, C, F, G, unpaired t-test (A, F, G: of AUC); ns: not significant, *p ≤ 0.05, **p ≤ 0.01, ***p ≤ 0.001, ****p ≤ 0.0001.

**Suppl. Figure 7:**
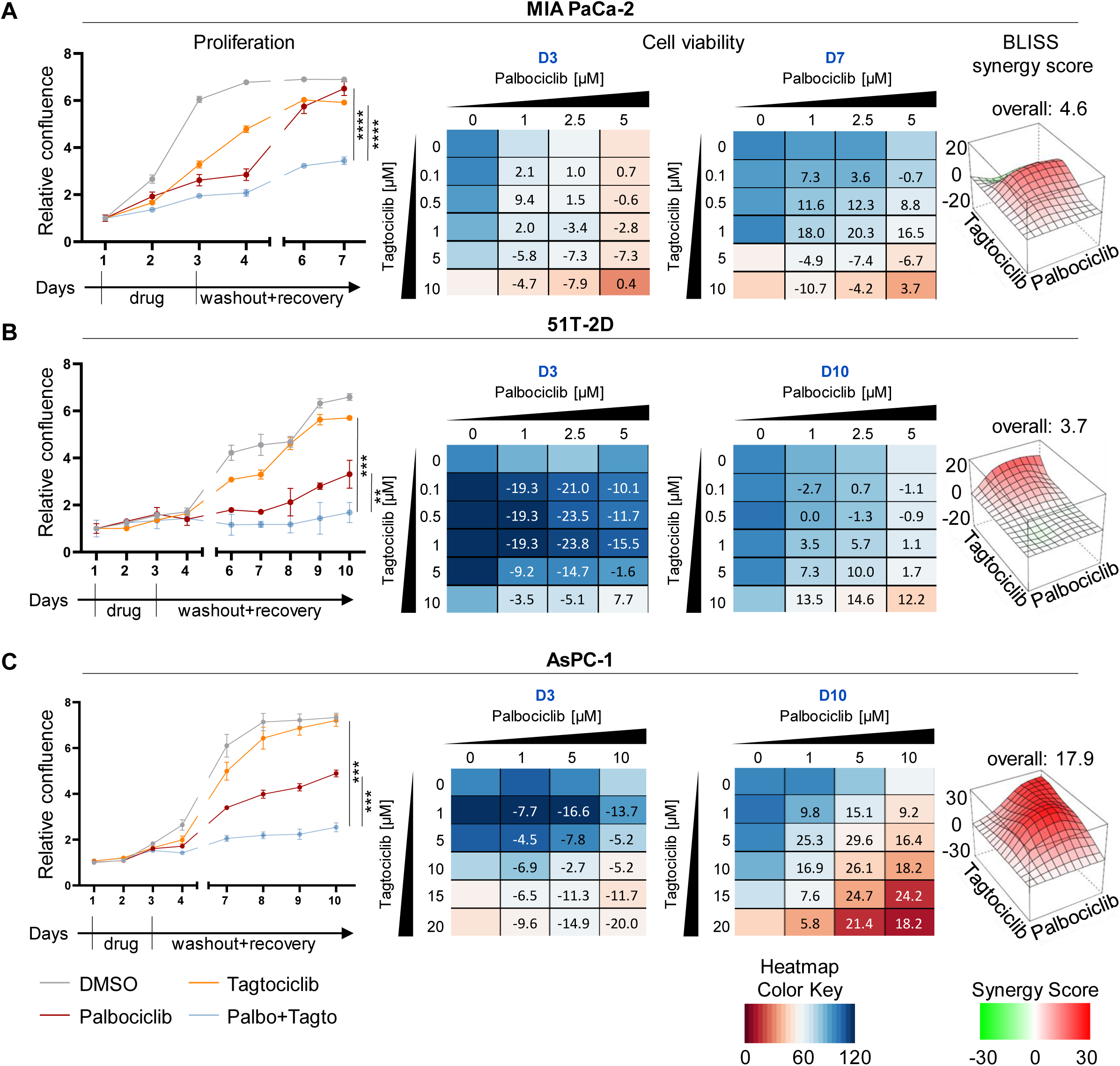
CDK2 inhibition can substitute for antagonizing KRAS in synergy with CDK4 inhibition. (A-C) All cells were treated and observed as in Suppl. Fig. 1, with the following specifics: (A) Proliferation of MIA PaCa-2 cells, 1 µM Tagtociclib, 5 µM Palbociclib, evaluation at D3 and D7. (B) Proliferation of 51T-2D cells, 5 µM Tagtociclib, 1 µM Palbociclib, D3 and D10. (C) Proliferation of AsPC-1 cells, 5 µM Tagtociclib, 5 µM Palbociclib, D3 and D10. Statistical analyses: A, B, C, unpaired t-test (A, B, C: of AUC); ns: not significant, *p ≤ 0.05, **p ≤ 0.01, ***p ≤ 0.001, ****p ≤ 0.0001.

## Notes

### Competing Interest Statement

The authors have declared no competing interest.

